# The More Unclear the Hearing, the Faster the Feeling: Speech Comprehension Changes Subjective Speech Rate

**DOI:** 10.1101/2023.10.17.562834

**Authors:** Liangjie Chen, Yangping Jin, Zhongshu Ge, Lingxi Lu, Liang Li

**Author notes:** **Corresponding Authors:** Liang Li, School of Psychological and Cognitive Sciences, Peking University, Lingxi Lu, Center for the Cognitive Science of Language, Beijing Language and Culture University.

## Abstract

Imagine being in a foreign country, surrounded by locals speaking a language entirely alien to you. Does their conversation seem to race like a machine gun? The perception of speech rate, as a facet of time perception, greatly impacts our daily lives. However, the factors influencing this perception remain poorly understood. Through a series of three experiments (with sample sizes of 31, 27, and 25 respectively), we discovered that speech comprehension significantly alters listeners’ perception of speech rate. As comprehensibility diminishes, listeners tend to overestimate the speech rate. Intriguingly, this effect is further influenced by the listener’s language proficiency. Specifically, the better the listener’s language proficiency, the more likely their perception of speech rate is influenced by the speech comprehension. Our study corroborates the notion that biases in time perception may be prevalent in our everyday communication. These subjective experiences could prompt listeners to modify their behavior or processing strategies.

## Statement of Relevance

Humans possess the innate ability to perceive the flow of time, despite lacking a specific sensory organ to receive time information. However, this perception seems to be easily fooled by various factors, resulting in no longer accurate. Indeed, we’ve discovered that such distortions in time perception are prevalent during speech processing. When speech becomes unclear or incomprehensible, listeners invariably overestimate the speech rate. Interestingly, this phenomenon is exclusive to native speakers or those with high language proficiency. Furthermore, the shift in subjective speech rate is not merely a simple time illusion. It can induce perceptual effects akin to those triggered by changes in objective speech rate. Our findings contribute to establishing a paradigm for time perception of complex auditory stimuli and pave the way for exploring more general mechanisms affecting time perception.

## The More Unclear the Hearing, the Faster the Feeling: Speech Comprehension Changes Subjective Speech Rate

“Unless we are told the reference body to which the statement of time refers, there is no meaning in a statement of the time of an event.” — Albert Einstein (1920, p. 32).

From the nicks marking the lunar calendar 10,000 years ago, to the shadows on the coronagraph 6,000 years ago, and now to the atomic clock with a precision of one ten-millionth of a second. humans have always yearned for an accurate depiction of time. Our brains have been demonstrated to possess the ability to finely delineate the dynamic temporal changes of both internal and external things (Harvey et al., 2020; Naghibi et al., 2023). In everyday discourse, listeners can identify temporal characteristics within speech. The processing of slow envelope fluctuations aids in our comprehension of semantics, while the rapid oscillations of fine structures enable speaker identification (Moore, 2019; Swaminathan et al., 2016). However, the question remains: can we truly perceive changes in temporal features as they occur?

Time perception can easily yield measurable distortions and illusions (Eagleman, 2008). An individual’s current attention, emotions, memory, and other cognitive processes, as well as the surrounding environment, will change our estimates of the duration or flow rate of a period (Block et al., 2010; Cui et al., 2023; Howard, 2018). Researchers typically employ filled or empty intervals for participants to reproduce or compare against a standard time template. However, these studies primarily focus on simple auditory stimuli, such as pure tones or noise segments, and overlook more complex or real-world acoustic objects like speech. IBut listeners are easily influenced by the auditory scene in which they are immersed when processing speech stimuli. For instance, being in a context with a fast speech rate can cause listeners to perceive the syllables they hear as long vowels or voiced consonants, and even perceive non-existent words (Baese-Berk et al., 2014; Maslowski et al., 2019; Wade & Holt, 2005). These results indicate that the temporal features of speech processing can be distorted.

Have you ever found it challenging to follow a speaker’s content in a noisy environment or when the language is unfamiliar, yet easily keep pace in a quiet setting or when the language is your native tongue, even if the speech rate remains constant in both scenarios? In studying time perception within the millisecond to second range of speech, researchers asked monolingual speakers to share their subjective experiences when processing different languages (Eveline, 1982), or to compare the duration of stimuli presented under varying degrees of word familiarity (Fernandes & Garcia-Marques, 2020). A recent study discovered that listeners overestimate the speech rate when in a non-native language context, irrespective of the actual speech rate (Bosker & Reinisch, 2017). Moreover, the encoding of familiar language content in memory exhibits greater processing efficiency and temporal information detection, leading to an expanded subjective duration (Jacoby & Dallas, 1981; Wänke & Hansen, 2015). These findings raise the speculation that the easier speech is to process, the slower it appears to sound.

We propose that speech comprehension influences the listener’s time perception. Speech comprehension refers to the process of understanding spoken language input. It involves recognizing speech signals and transforming them into meaningful information, which includes acoustic analysis and speech recognition to achieve comprehensible and interpretable semantic perception (McGettigan et al., 2012; Price, 2003). When speech comprehension is compromised, such as in noisy environments or with difficult-to-understand language, the amount of semantic information that listeners can extract decreases. Theories of time perception, whether it’s the encoding efficiency hypothesis (Eagleman & Pariyadath, 2009), the processing principle (Matthews & Meck, 2016), or the pacemaker-accumulator model where arousal and attention affect the number of pulses (Grondin, 2010), all imply that an increase in perceived time information leads to an increase in an individual’s duration estimation. Variations in speech comprehension can cause a deviation in the amount of time information perceived by the listener, potentially leading to changes in time perception. We hypothesize that when speech comprehension decreases, the reduction in decodable information from speech results in time compression, perceived as a subjective increase in speech rate.

Our goal is to verify the influence of speech comprehension on the subjective speech rate. Prior research has established that speech comprehension relies on the spectral content of the speech signal and the integrity of the temporal envelope (Ahissar et al., 2001). In Study 1, we manipulated the spectral features of clear speech to decrease its comprehensibility, aiming to observe the subjective speech rate under varying levels of comprehensibility. In Study 2, to avoid additional interference from changes in acoustic features, we constructed syntactically congruent simple sentences and random syllable sequences devoid of syntax to manipulate comprehensibility levels. We combined this with an implicit speech rate task to detect differences in speech rate perception.. In the final Study 3, we recruited a non-native Chinese group with different level of Chinese language proficiency to replicate the experiment in Study 2. We wondered if the bias in subjective speech rate induced by speech comprehension is modulated by the participants’ language proficiency, further substantiating the impact of speech comprehension on speech rate perception.

## Study 1: Listeners judge the speech rate in noise-vocoder speech

### Method

#### Participants

We enlisted 34 adult participants (average age: *M* = 22.355 years, *SD* = 2.801, 21 females) from Peking University. All participants were native Chinese speakers, born and currently living in China, and participated in this study for course credit or monetary compensation. Written informed consent was obtained before the experiment started. None of the participants reported any hearing or vision difficulties.

The sample size of this study was determined on the previous studies about judging speech rate (Pfitzinger & Tamashima, 2006; Plug & Smith, 2021). In study by Pfitzinger and Tamashima (2006), with 20 participants in each group, they found effect sizes (*η*^2^ = .267 ∼ .324) for the main effects of different languages in the speech rate judgment task. Another study (Plug & Smith, 2021) was recruited 34 participants and was found a robust effect size (*η*^2^ = .810) for judging speech rate when fixing the constituent duration. In our experiment, all stimuli used are spectral attenuation speech, implying a higher processing difficulty compared to the clean speech used in previous studies. Therefore, we recruited 34 participants to ensure a sufficient effect size. Three participants (2 females) were excluded: Two participants used a single response option across all trials (e.g., always selecting faster option). One of the participants abandoned the experiment. The remaining 31 participants underwent formal data analysis.

##### Stimuli

All sentences were sourced from the Comprehensive Language Knowledge Base (CLKB, https://klcl.pku.edu.cn) established by the Key Laboratory of Computational linguistics of Peking University. We selected a total of 100 sentences, each a 12-word statement with a “subject + predicate + object” syntactic structure. These clean speeches were recorded by a 23-year-old female native Chinese speaker. The durations of speeches ranged from 2.92 to 3.14 seconds (approximately 3.7 Hz, see Fig. S1). All speeches were subjectively rated for comprehension and speech rate to ensure consistency (see Fig. S2).

Eighty sentences were chosen for speed ratings in the formal experiment and underwent spectral attenuation operations to produce noise-vocoder speech. The remaining 20 sentences served as standard speed templates. The noise-vocoder speech was created by dividing the clean speech signal into logarithmically spaced frequency bands (80 - 8820Hz). The amplitude envelope of each frequency band was then extracted and used to modulate the temporal fine structure of speech-spectrum noise (with the same power spectrum as the clean speech) in the same frequency band. All bands were then recombined (the details in Supplementary material). Through an intelligibility test, the frequency bands of the noise-vocoder speech were selected near the 50% intelligibility threshold for each participant.

The Time-Domain Pitch Synchronous Overlap and Add (TD-PSOLA) technology in Praat (Boersma, 2001) was used to linearly compress or expand the formal experimental speech, with the rate factor set to 0.8, 0.9, 1, 1.1, 1.2, totaling 5 levels (1 represents the original speech rate). These speech recordings were normalized for the root-mean-square intensity to broadly equalize the perceived loudness. All stimuli were always presented binaurally via headphones (Sennheiser HD569, Sennheiser Corporation, Old Lyme, CT).

#### Design and Procedure

All experiments were run in a sound-attenuated and dimly lit room. Firstly, participants commenced the speech familiarity task, in which they were asked to listen to 40 clean speeches randomly selected from 80 clean speeches (see Fig. 1). The listeners had the freedom to repeat these speeches as many times as needed, until they were familiar with them. The noise-vocoder of these sentences would serve as high-comprehension (HC) condition in formal experiments. The remaining 40 noise-vocoder speeches were treated as low-comprehension (LC) conditions.

**Fig. 1.**
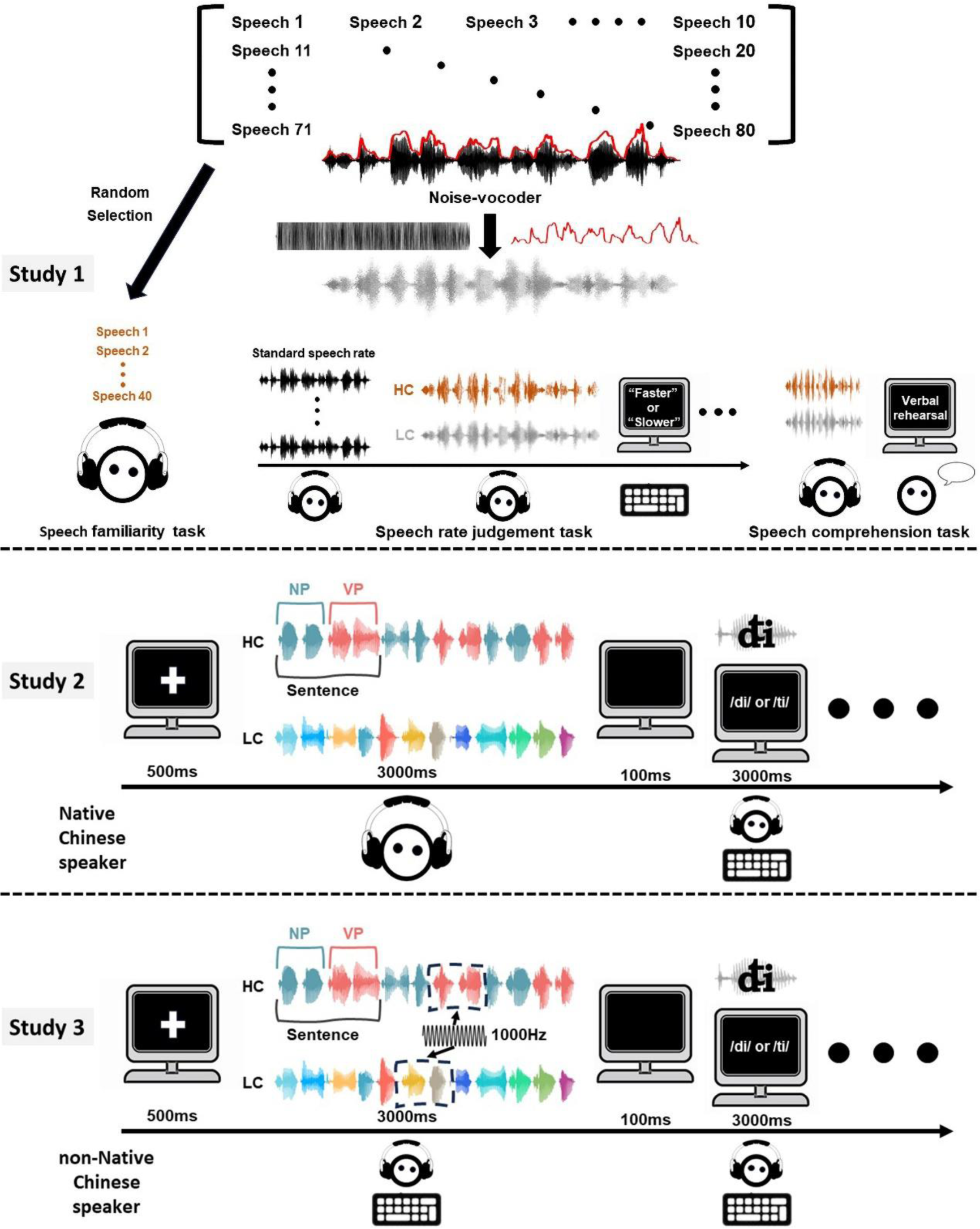
Illustration of Experimental Procedure (Not Drawn to Scale) *Note.* Top: In Study 1, we initially extracted 40 speeches from the selected 80 for the speech familiarity task. Subsequently, participants undertook the speech rate judgement task. Finally, participants completed a speech comprehension test to determine whether the familiarity task improved comprehensibility of speech. Middle: In Study 2, participants performed an implicit speech rate task. Participants’ task was to classify the last appearing morphing syllable. Bottom: In Study 3, we replicated the experiment from Study 2 among non-native Chinese speakers. Additionally, we included some probe stimuli with pure tone segments (1000Hz) to ensure participants’ attention.

Secondly, participants started the speech rate judgement task, which was consisted of four blocks. At the beginning of each block, participants were informed that they would be hearing five clean speeches as standard speech rate. These speeches are derived from the remaining 20 sentences and do not appear in all blocks. Five speech rates were used: 0.8, 0.9, 1,1.1. 1.2. These speech rates were factorially combined with the two types of noise-vocoder speech (i.e., HC and LC condition), resulting in 10 conditions. In each block, every condition is rendered 10 times and the order of trials was randomized. Each block was interleaved with a 2 min break.

Each trial started with a fixation cross which was presented for 500 ms. Then, when the noise-vocoder speech was over, participants were asked to judge whether the speech rate of noise-vocoder speech faster or slower than the standard speech. Participants were encouraged to swiftly provide their ratings, guided by their initial reactions rather than overthinking their choices. And they were stressed to avoid using beats or counting to help them decide. The response window was presented for a maximum of 3 s or until participants responded. Trials were spaced apart by an inter-stimulus interval (ISI) ranging from 1000 to 1500 milliseconds, with the duration being randomly allocated on an individual trial basis.

Thirdly, noise-vocoder speeches with the factor of speech rate set to 1 in the speech rate judgement task were used for speech comprehension test. Participants were instructed to verbally repeat the entire sentence as much as possible when the speech ended in each trial. The accuracy calculated the number of correctly recognized phrase of the object phrase (Wang et al., 2021). The difference in accuracy between HC and LC speech was shown in the supplementary materials (see Fig. S3).

#### Statistical analysis

For each participant, we calculated the proportion of responses indicating “slower” speech rate as a function of the different rate factor, separately for both the HC and LC conditions. We modeled participants’ psychometric curves using a Weibull function, employing a maximum likelihood procedure (Fig. 2). Then, we extracted the Point of Subjective Equivalence (PSE) and Difference Limen (DL, computed as the difference between the 25% and 75% performance levels for each participant) for everyone. The PSE corresponds to the subjective speech rate that is perceived as equivalent to the standard speech rate, with a smaller PSE value indicating a relative underestimation of the speech rate. The DL reflects participants’ discrimination sensitivity, with smaller values signifying higher sensitivity.

**Fig. 2.**
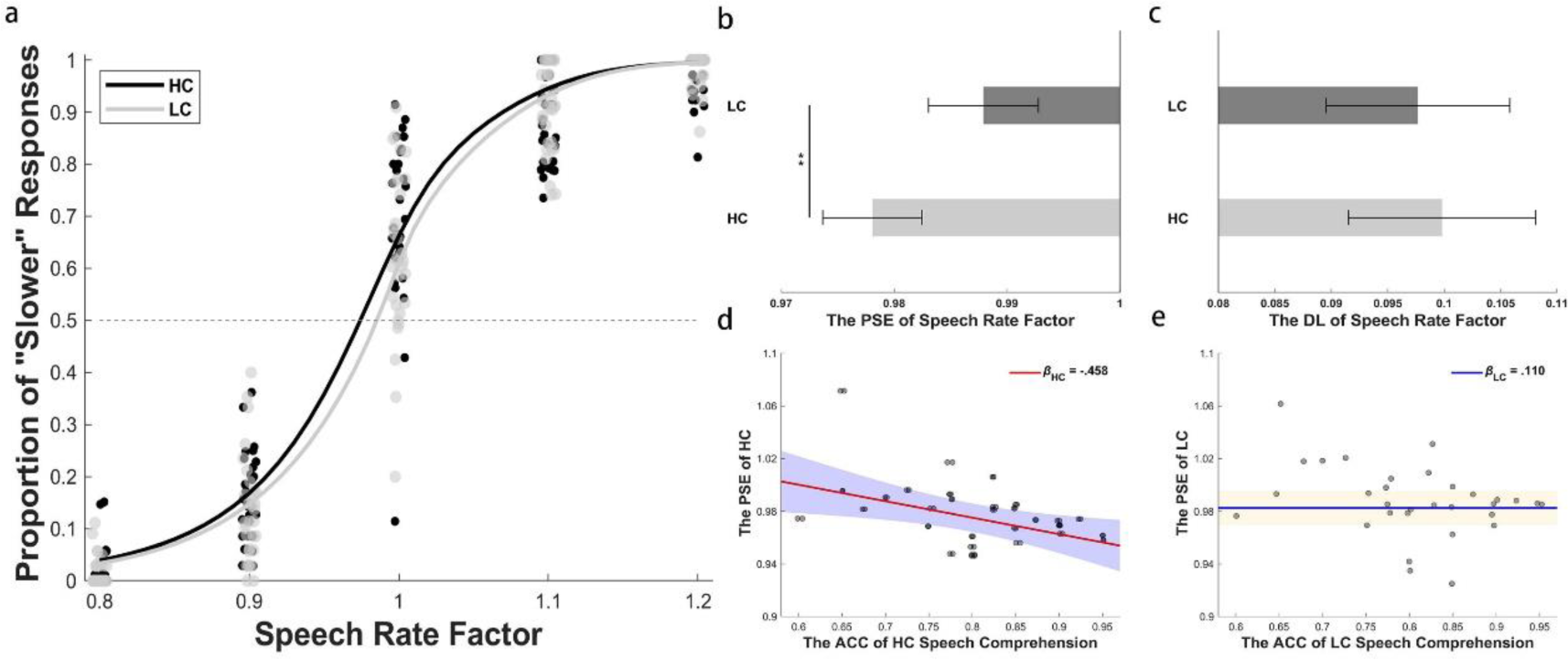
The results of point of subjective equality (PSE) and difference limen (DL). *Note.* (a): The mean psychometric curves were fitted to the grand average data points. Each solid circle (high comprehension: dark; low comprehension: light gray) represents the proportion of “shorter” responses for each individual participant. (b): Mean PSE derived from Weibull psychometric fits on a per individual in HC and LC condition. The mean PSE in HC (light gray) was significantly smaller than the PSE in LC (dark gray). (c): Mean DL derived from Weibull psychometric fits on a per individual in HC and LC condition. The mean DL in HC (light gray) did not significantly differ from the mean DL in LC (dark gray). (d) and (e): Scatterplots (with best-fitting hierarchical regression lines) showing the relationship between the ACC of HC/LC in the speech comprehension test and the PSE of HC/LC in the speech rate judgement task. Results are shown negative relationship. Bars are standard error. Filled areas represent 95% confidence intervals. **: *p* < .001.

A paired t-test was performed on the parameters PSE and DL between HC and LC condition. Hierarchical multiple regression analysis was then used to assess the speech comprehension to predict the PSE, while controlling for the effects of age and gender. And the enter method was used. In step 1, the PSE score was the dependent variable, with age and gender serving as independent variables. In step 2, we added the ACC of speech comprehension task to the model from step 1. Preliminary analyses were performed to ensure no violation of the assumptions of independence and multicollinearity (all tolerance values were greater than .9). We also calculated Bayes factors (*BF*_10_) for the null hypothesis with frequentist inferential statistics. All statistical analyses were performed in JASP 0. 17.3 software (2023).

### Results

#### Effect of the speech comprehension on PSE and DL

Figure 2a presents the percentage of “slower” responses for all participants, modeled as a Weibull function of five speech rate factors under both HC and LC conditions. Prior to the paired t-test, we analyzed the data distributions using the Shapiro-Wilk test for normality (*p* = .361).

As expected, a paired t test using the PSE as dependent variable revealed the effect of speech comprehension. The result showed that the grand average PSE in the HC (*M* = .978, *SD* = .024) significantly smaller (*t* = −3.335, *df* = 30, *p* = .002, Cohen’s *d* = −.599, see Fig. 2b) than the average PSE in the LC (*M* = .988, *SD* = .0.27). This suggests that high comprehension corresponds to more “slower” responses for noise-vocoder speech following a familiarity task. Fig. 2c displays the DL in HC and LC. The results revealed no significant differences between HC and LC (HC: *M* = .100, *SD* = .046; LC: *M* = .098, *SD* = .045; *t* = .374, *df* = 30, *p* = .711, Cohen’s *d* = .067, *BF*_10_ = .204, anecdotal evidence for the null hypothesis).

The decrease in PSE indicates a tendency for listeners to perceive the speech rate as slower. These psychophysical results suggest that speech comprehension influences the participant’s perception of speech rate. Compared to the standard speech rate, noise-vocoder speeches with low comprehension may sound faster than those with high comprehension. Surprisingly, through the one sample t test, we find that PSE under both conditions (HC: *t* = −4.990, *df* = 30, *p* < .001, Cohen’s *d* = −.896; LC: *t* = −2.487, *df* = 30, *p* = .019, Cohen’s *d* = −.447) is significantly less than 1, indicating the participant thought both conditions were faster than the standard rate (see Fig. S4). We suspect this is due to the operation of the noise-vocoder. Even if part of the speech has undergone familiarization training, all the noise-vocoder speech chosen for the formal experiment are close to the 50% comprehensibility threshold, which is a relatively low level. Consequently, there is a general tendency to overestimate speech rate.

#### Speech comprehension predicts the PSE

Given that low comprehension noise-vocoder speech led to an overestimation of speech rate, we investigated whether the ACC in speech comprehension test could predict the variability of the PSE. Correlation analyses revealed a negative relationship between the ACC of HC speech and the PSE of the HC condition (*r* = −.453, *p* = .011). However, no significant correlation was found between the ACC of LC speech and the PSE of the LC condition (*r* = .138, *p* = .458). Following the methodology described, Model 1 results showed that age and gender accounted for a variance (*R^2^*) of .044 in the PSE (adjusted (*R^2^*) = −.024), which was not significant (*F* (2, 30) = .650, *p* = .530). After entry of the ACC of HC speech in Model 2, the total variance explained by the model was 0.25 (adjusted *R^2^*= .167), which was significant (*F* (3, 30) = 3.004, *p* = .048). In Model 2, the standardized coefficient of the ACC was significantly (*β* = −.458, *t* = −2.723, *p* = .011, see Fig. 2d), while other standardized coefficients were not (Age: *β* = −.055, *t* = −.322, *p* = .750; Gender: *β* = −.195, *t* = −1.142, *p* = .263). The same analysis was conducted for the ACC of LC speech and the PSE of LC condition. However, no significant results were found. In Model 2, the ACC of LC speech comprehension only explained 8.7% of the variance (adjusted *R^2^*= −.014) and the standardized coefficient was not significant (ACC: *β* = .110, *t* = −.592, *p* = .559, see Fig. 2e). In summary, these results suggest that the PSE of the HC condition was associated with speech comprehension. Furthermore, the findings indicate that as speech comprehension decreases, the subjective perception of speech rate tends to faster.

Overall, these results align with an overestimation of speech rate in the low comprehension condition. However, manipulating the comprehensibility of noise-vocoder speech through familiarity may introduce judgment bias based on memory. To further explore the relationship between speech comprehension and the perception of speech rate, we conducted Study 2, utilizing clean speech and an implicit rate perception task (Bosker & Reinisch, 2017). Additionally, we employed the drift diffusion model (DDM) to demonstrate that the bias in speech rate perception is not merely a categorization strategy but can also produce the same perceptual effects as the objective speech rate.

## Study 2: Subjective speech rate was assessed by implicit speech rate task

In this study, we employed the implicit rate perception task to assess the perception of speech rate. The speech rate of a preceding speech can influence the perception of subsequent vowels or consonants in a syllable (Bosker & Reinisch, 2017; Wade & Holt, 2005). Specifically, when listeners are exposed to fast speech, the perception of the following syllable is biased towards a longer vowel or voice onset time of the consonant. This is the rate normalization effect. Thus, we can infer the listener’s subjective speech rate from the result of syllable classification.

### Method

#### Participants

We recruited 27 adult participants (age: *M* = 22.593 years, *SD* = 2.735; 14 females) from the Peking university. All participants took part in this study for course credit or payment and did not participate in Study 1. Each participant provided written informed consent. None had any self-reported hearing or vision difficulties.

The sample size of this study was determined on the previous studies about the implicit rate perception task (Bosker & Reinisch, 2017). In their studies, with 20 to 27 participants, they found medium effect sizes (*η^2^* = .018 - .068) for the main effects of different speech rate. Because our study is adapted from their task, the simple size of 27 is selected to ensure the sufficient effect size.

##### Stimuli

We manipulated the level of speech comprehension by constructing simple and random sentences. Simple sentences, serving as high comprehension (HC) speech, are composed of noun phrase (NP) and verb phrase (VP) that adhere to Chinese syntax, i.e., “subject + predicate”. We selected a total of 50 simple sentences from the study by Ding et al. (2016) with each phrase consisting of two syllables (each has 250 ms). Random sentences (total 50 sequences), serving as low comprehension (LC) speech, are sequences generated by randomly recombining the individual syllable from all simple sentences and manually checked to exclude syntactic or semantic material.

The syllables used in this study are /di/-/ti/, and the pronunciation rate of the syllables aligns with the speech rate of above syllable material (/di/ lasts for 249ms, /ti/ lasts for 289ms). We employed the “progressive cutback and replacement” method to create the morphing syllables (Winn, 2020). Prior to the formal experiment, we determined each participant’s classification threshold for the syllable continuum. Based on this threshold, we selected syllables that were closest to the 25%, 50%, and 75% subjective classification points as target morphing syllable for the subsequent experiments. For more details, please refer to the supplementary materials.

#### Design and Procedure

Each trial started with the presentation of a fixation cross with 500ms (see Fig. 1). Following this, the speech was played, and 100ms after the sentence ended, a target morphing syllable was played. Meanwhile, participants were asked to judge whether they heard /di/ or /ti/. After their response (or timeout after 3 s), the screen was replaced by an empty screen for 500 to 1000ms, after which the next trial was performed. This study is a within-subjects design with 6 conditions, comprising 3 morphing levels (25%, 50%, and 75%) and 2 speech comprehension levels (HC and LC). Each condition consisted of 40 trials, totaling 240 trials. All trials were randomly distributed across 4 blocks, with a minimum rest period of 2 minutes between blocks. During the experiment, the participants’ responses and reaction times for each target syllable were recorded.

##### Statistical analysis

The perception of speech rate differences was examined by comparing the proportion of /ti/ responses to morphing syllables under simple speeches (HC) and random syllable sequences (LC).

Trials where no judgment was made within 3 seconds were excluded, accounting for less than 1% of the total data. A repeated-measures ANOVA was conducted with Holm-Bonferroni corrected post hoc tests for significant differences. All statistical analyses were performed using JASP 0.17.3 software (2023).

When listeners exhibit a bias in syllable classification between HC and LC conditions, two scenarios are possible. The first is that comprehension leads to differences in the perception of speech rate, causing a bias classification (rate normalization effect). The second scenario is that participants classify syllables based on the type of speech. For instance, when hearing a simple sentence, they tend to classify the syllable as /ti/. We used the DDM to identify the specific scenario. In the first scenario, the bias in classification induced by the speech rate should occur in the syllable encoding phase and provide no clues about the upcoming syllable. Therefore, the drift rate, which characterizes the accumulation of noisy evidence over time, may vary across conditions. However, the starting point, which can be conceptualized as a statistical prior and characterizes the initial bias before any information about choice attributes becomes available, should remain consistent (Kloosterman et al., 2019; Ratcliff & McKoon, 2008). In the second scenario, the opposite is true.

In the null model, all parameters were fixed across conditions. The target model allowed the drift rate to vary with all conditions, and the starting point to change with HC and LC conditions, while other parameters remained fixed across conditions. All models were fitted to stimulus-coded data (i.e., the upper and lower boundaries corresponded to /ti/ and /di/ responses) using hierarchical Bayesian estimation. This method captures group-level commonalities and estimates each participant’s parameter values (Ratcliff et al., 2016). The HDDM toolbox (v 0.9.7, Wiecki et al., 2013) was used in Python. We used Markov chain Monte Carlo sampling to estimate the posterior distribution of each model and ran 4 chains with 5000 samples, discarding the first 500 samples as burn-in. Convergence was assessed by visual inspection (the trace and autocorrelation) and the Gelman-Rubin R^ statistic. The range of R^ statistic values across all group parameters in each model was 0.99 to 1.01, suggesting good convergence. Goodness of fit was visually inspected with a posterior predictive check (see Fig. S5). The Deviance Information Criterion (DIC) was used for model comparison. Bayesian hypothesis testing was performed by analyzing the probability mass of the parameter at the group level (e.g., examining the overlap of the posterior distributions of parameters).

### Results

#### Effect of the speech comprehension on the classification of morphing syllable

We examined whether different speech comprehensions would lead to bias in the syllable classification. The results revealed a significant main effect of speech comprehension (*F*(1,26) = 24.295, *p* < .001, *η^2^*= .015, see Fig. 3a). Post-hoc comparisons revealed that the proportion of choosing /ti/ was higher in the LC condition (*M* = .653, *SE* = .025) compared to the HC condition (*M* = .585, *SE* = .019, *t* = 4.929, *p* < .0001). Base to the rate normalization effect, when listeners are exposed to faster speech prior to a morphing syllable, they tend to perceive the VOT of consonant in this syllable as longer (Wade & Holt, 2005), such as /ti/. Thus, our results were consistent with the subjective speech rate obtained in Study 1, where listeners tended to perceive the speech in the LC condition as faster compared to the HC condition. The main effect of morphing level was significant (*F*(2, 52) = 88.922, *p* < .001, *η^2^* = .736), indicating that increasing the duration of VOT induced all listeners to report more /t/ responses.

**Fig. 3.**
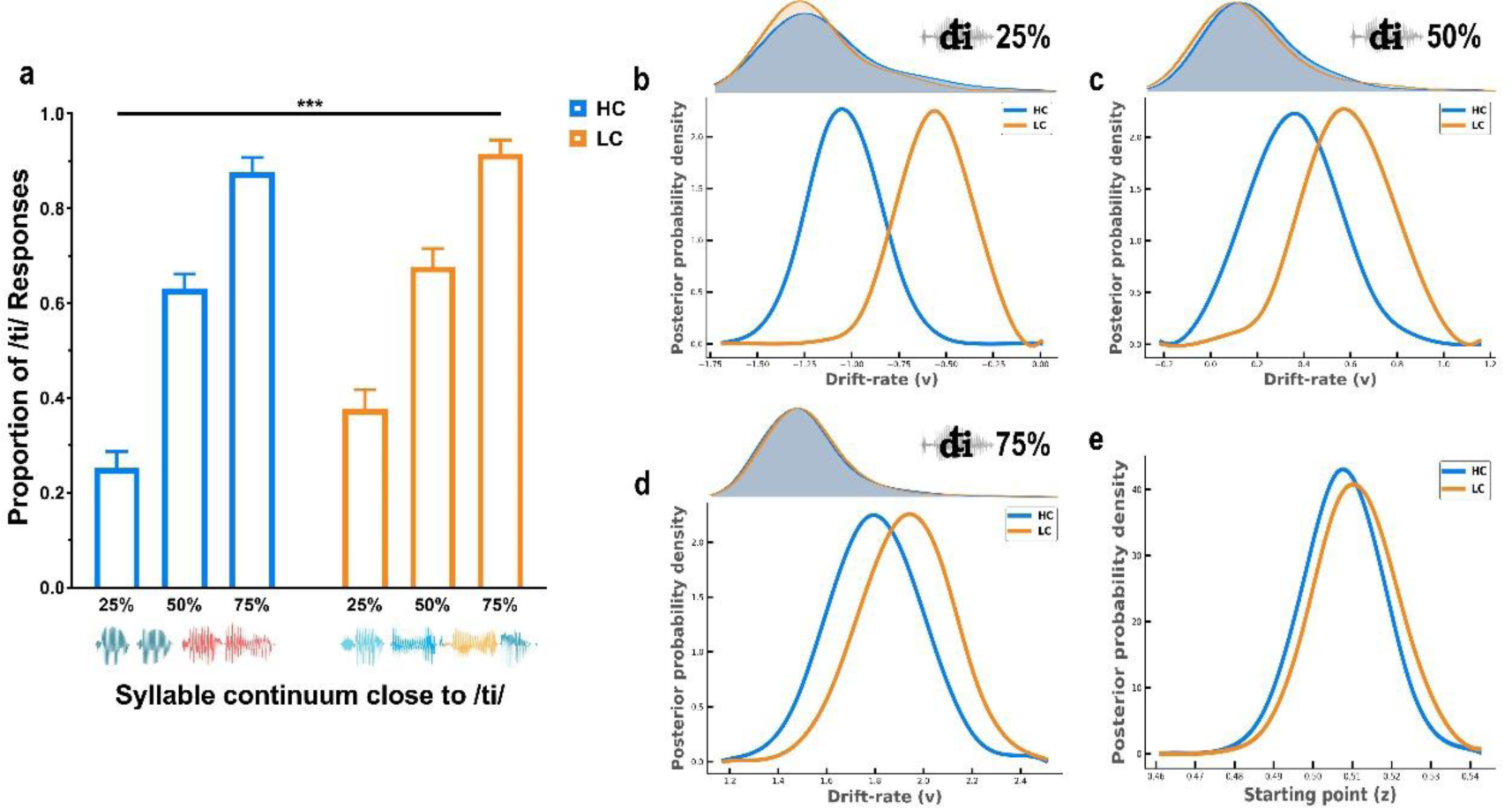
The result of the classification of morphing syllable and DDM. *Note.* (a): The proportion of participants choosing /ti/ under different conditions. Under the low comprehension (LC) condition, the proportion of participants classifying the morphing syllable as /ti/ is larger, compared to high comprehension (HC) condition. This difference exists at all morphing levels. (b) – (d): The upper part shows the response time distribution of choosing /ti/ at each morphing level. The lower part shows the posterior probability distribution of the drift rate under different comprehension conditions. (e): The posterior probability distribution of the starting point under different comprehension conditions. ***: *p* < .001.

Importantly, the interaction between the two factors was significant (*F*(2,52)=3.235, *p* = .047, *η^2^* = .002). Through simple effect analysis, we found that the rate normalization effect induced by speech comprehension consistently appeared across different levels of morphing syllable: In 25%, *M*_LC_ (.363) > *M*_HC_ (.262), *p* < .001; In 50%, *M*_LC_ (.682) > *M*_HC_ (.619), *p* = .003; In 75%, *M*_LC_ (.914) > *M*_HC_ (.874), *p* = .041. Thus, the above results revealed that the LC condition corresponded to higher /ti/ responses or a perception of faster speech across all morphing levels of the syllable, compared to the HC condition.

#### Speech comprehension changes the drift rate, not the starting point

Model comparison favored the target model (ΔDIC relative to null model: = −2699.929). Figure 3b, c and d showed the group mean posterior estimates of the drift rates for each condition and supported the first scenario: the drift rate changes, but the starting point does not differ. Drift rate in the LC condition was generally larger than in the HC condition (25%: posterior *p*_LC > HC_ = .966; 50%: posterior *p*_LC > HC_ = .808; 75%: posterior *p*_LC > HC_ = .675).

Higher drift rates in this study favoring one option over the other generate faster reaction time for the favored option. The RT distribution also showed a larger proportion of the leading edge in the low comprehension (LC) condition. These findings indicated that an overall accumulation bias towards the bound for longer VOT, i.e., /ti/. It is worth noting that there was little difference in the leading edge of RT at the 75% level of morphing syllables, and the posterior probability was close to 0.5. We think that the 75% morphing syllable is perceived to be very close to /ti/, which means it is less affected by the rate normalization effect. Additionally, we analyzed the differences in the drift criterion (*dc*) under the HC and LC conditions (refer to supplementary materials). The results also confirmed that changes in speech comprehension alter the tendency of evidence collection, with a larger dc observed in the LC condition (LC > HC: posterior *p* = .968, see Fig. S8).

We found no evidence for differences in the starting point between conditions (posterior *p*_LC > HC_ = .582, see Fig. 3e). The starting point refers to an automatic pre-decisional response bias. The result indicates that participants do not have any classification bias before hearing morphing syllables. Thus, they do not classify syllables by speech type.

#### Correlation between performance on the classification of morphing syllable and drift rate

We conducted Pearson correlation analyses between the Classification Bias Index (CBI) for each level of morphing syllable and drift rate. The CBI is calculated as the proportion of /ti/ responses minus the individual morphing level, divided by the individual morphing level. If a positive correlation emerges, it will indicate that listeners align more closely to the DDM in syllable classification tasks. This means their responses would be faster and exhibit a stronger bias towards longer VOT of /ti/.

Drift-rate parameters were estimated for each participant in the target model. We found that participants with larger BI exhibited higher drift rate in the 25% and 50% levels of morphing syllables (25% in HC: *r* = .773, *p* < .001, Fisher’s *z* = 1.029; 25% in LC: *r* = .566, *p* = .003, Fisher’s *z* = .652, see Fig. 4ab; 50% in HC: *r* = .658, *p* = .0002, Fisher’s *z* = .789; 50% in LC: *r* = .761, *p* < .001,

**Fig. 4.**
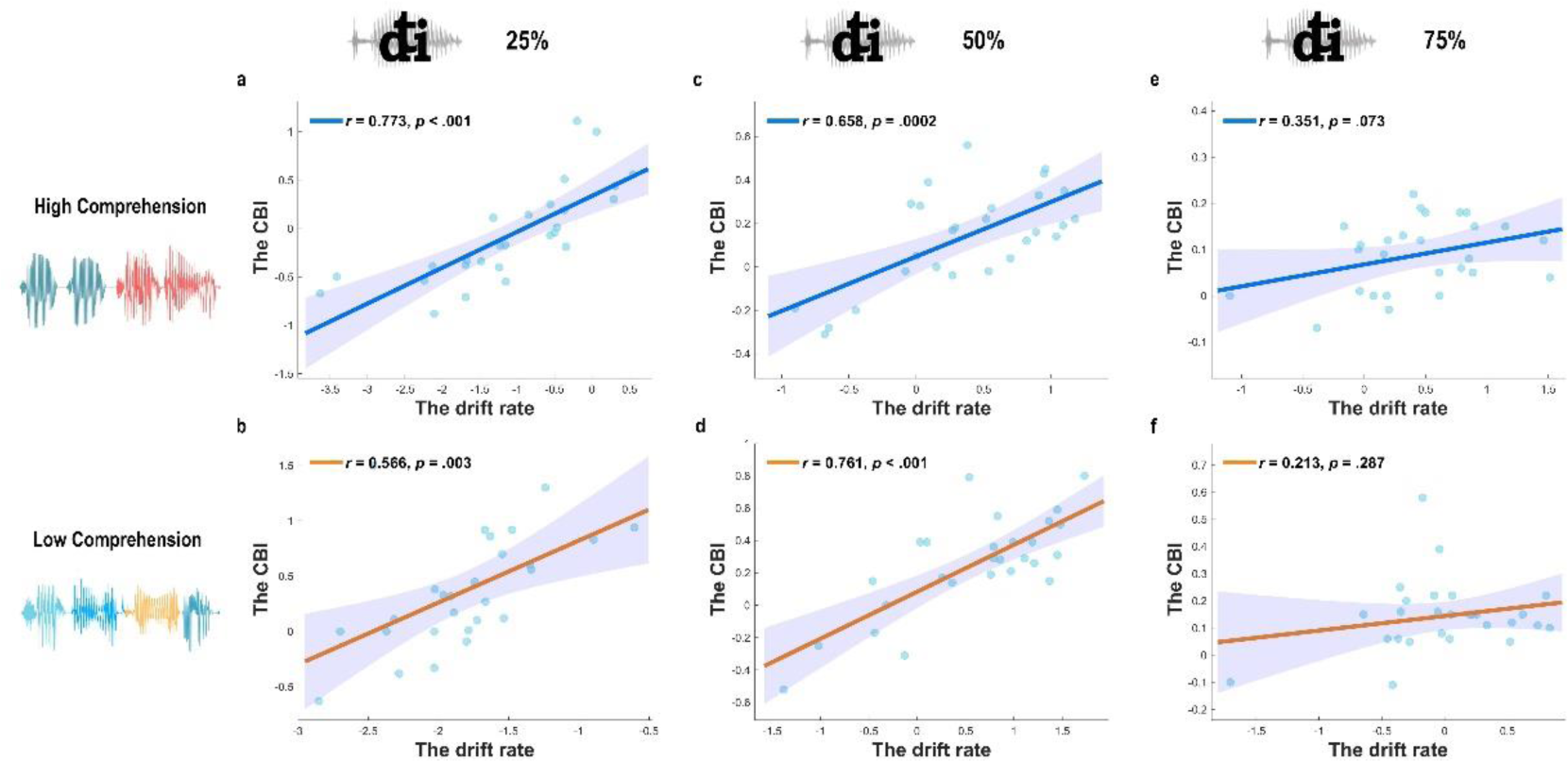
Correlation between the Classification Bias Index (CBI) and the drift rate across all condition. *Note.* (a), (c) and (e) are the correlation results under the high comprehension (HC) condition. (b), (d) and (f) are the correlation results under the low comprehension (LC) condition. The solid line represents the best-fitting regression. The shaded region reflects the 95% confidence interval.

Fisher’s *z* = 1.000, see Fig. 4cd). However, no correlation was found at the 75% level (75% in HC: *r* = .351, *p* = .073, Fisher’s *z* = .204; *BF*_10_ = 1.107; 75% in LC: *r* = .213, *p* = .287, Fisher’s *z* =.204, *BF*_10_ = .410, see Fig. 4ef).

The results of syllable classification reinforced the conclusions of Study 1: speech comprehension can modify a listener’s perception of speech rate. Furthermore, the DDM results demonstrated that this shift in perception is not just an auditory illusion. Essentially, it reflects the impacts of real speech rate differences, affecting the perception of subsequent stimuli through the rate normalization effect.

## Study 3: Language proficiency moderates the effect of speech comprehension on implicit speech rate

In this study, non-native Chinese speakers were recruited, and the experiment from Study 2 were repeated. They have different levels of Chinese language proficiency (CLP). Individuals with lower CLP are unable to process the semantics of simple sentences correctly and effectively, which makes the differences in speech comprehension between the two stimuli impossible to express. If speech comprehension does indeed alter the perception of speech rate, then no syllable classification bias should be detected in these groups. Thus, we hypothesize that the level of CLP among individuals can modulate the magnitude of syllable classification bias, with higher proficiency groups showing a stronger rate normalization effect.

### Method

#### Participants

The sample size was consistent with that of Study 2. We recruited 27 non-native Chinese adult participants (age: *M* = 22.880 years, *SD* = 3.370; 17 females) from the Beijing Language and Culture University and the Peking university. All participants took part in this study for payment. The level of Hanyu Shuiping Kaoshi (HSK, i.e., Chinese Proficiency test), a famous language test for non-native speakers’ abilities of using Chinese, is used to measure their CLP. We primarily aimed to investigate whether the Chinese language proficiency of participants would affect the role of speech comprehension in speech rate perception. Therefore, we did not strictly control for the participants’ native language backgrounds. Two participants abandoned the experiment. The remaining 25 participants underwent formal data analysis. All the participants had normal hearing and vision abilities, according to their self-reports.

#### Design and Procedure

All stimuli and procedure were mostly consistent with Study 2. To maintain participants’ attention to the speech, we added 15 probe trials in each block. During these trials, a 1000Hz pure tone with a duration of 500ms was randomly inserted into the speech fragments (see Fig. 1). Participants were instructed to make a keypress response (i.e., “SPACE”) as soon as they heard the pure tone.

#### Statistical analysis

First, the repeated-measures ANOVA and HDDM were used to replicate the analysis in Study 2, aiming to detect whether speech comprehension in non-native Chinese participants produces a classification bias in syllable perception. Then, to investigate whether Chinese language proficiency moderated the effect of speech comprehension on speech rate perception, we conducted moderation analyses using the PROCESS macro in SPSS 24.0 (Hayes, 2013). We evaluated the moderating effect through a simple mediation model (Model 1 in PROCESS Macro) We used the difference in response proportion for /ti/ between the HC and LC condition as the dependent variable. The difference in drift rate between two conditions was used as the independent variable, with Chinese language proficiency as the moderating variable. All continuous variables were standardized before the analysis. Gender and age were controlled in all analyses.

### Results

#### Non-native Chinese speakers did not exhibit the rate normalization effect

Firstly, like Study 2, the main effect of morphing level is significant (*F*(2, 48) = 117.080, *p* < .001, *η^2^* = .808), indicating that the classification of the syllables /di/ and /ti/ by non-native Chinese speakers is also influenced by the length of VOT. As expected, we did not observe a significant main effect of speech comprehension on syllable classification (*F*(1,24) = .006, *p* = .940, *η^2^*= 2.594*10^-6^, *BF*_10_ = .230, see Fig. 5a). Furthermore, the interaction between the two factors is not significant (*F*(2, 48) = .629, *p* = .537, *η^2^* = .0004, *BF*_10_ = .179). These results showed that there was no bias in morphing syllable classification between the HC and LC condition. Secondly, the DDM results also corroborate the above conclusion. The posterior distribution did not reveal evidence of differences in the drift rate (25%: posterior *p*_LC > HC_ = .460; 50%: posterior *p*_LC > HC_ = .536; 75%: posterior *p*_LC > HC_ = .493, see Fig. 5b - f) and starting point (*p*_LC > HC_ = .501). This suggests that regardless of the type of speech presented before the morphing syllable, it will not induce a bias in evidence accumulation(i.e., v) or pre-decisional response (i.e., z).

**Fig. 5.**
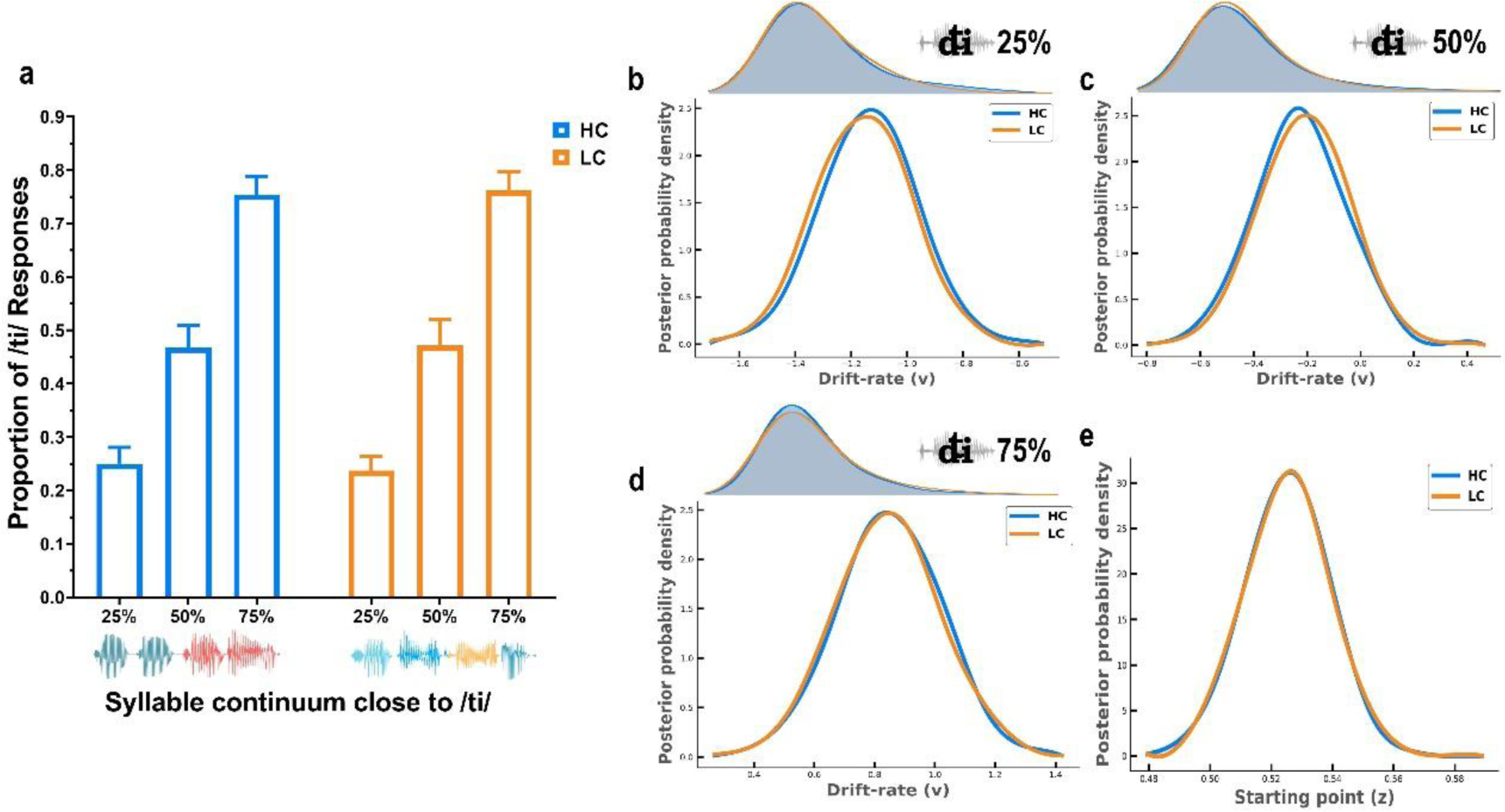
The result of the classification of morphing syllable and DDM in non-native Chinese speakers. *Note.* (a): The proportion of participants choosing /ti/ under different conditions. There is no difference between high comprehension (HC) and low comprehension (LC)condition exists at all morphing levels. (b) – (d): The upper part shows the response time distribution of choosing /ti/ at each morphing level. The lower part shows the posterior probability distribution of the drift rate under different comprehension conditions. (e): The posterior probability distribution of the starting point under different comprehension conditions.

The above results are based on group-level statistical analysis. We are intrigued to know whether non-native Chinese participants would show a bias in syllable classification as their Chinese language proficiency improves. In the next analysis, we will examine this hypothesis through a moderation effect analysis.

#### Chinese language proficiency moderates the effect of speech comprehension on the bias in syllable classification

We defined those with an HSK level above 5 as the high Chinese language proficiency group, and those with a level below 5 as the beginner-medium (low) Chinese language proficiency group (Lu et al., 2023). This dichotomy does not represent an absolute difference in Chinese language proficiency between groups, but it helps us detect potential moderation effect.

The moderation analysis examined the potential conditional effects of CLP on the direct process between the drift rate and the bias classification of morphing syllable across three morphing syllable conditions. In 50% morphing syllable, the difference in drift rate × CLP interaction had a significant effect on the difference in response proportion (*B* = .103, *t* = 2.562, *p* = .019, *R^2^* change = .069, in Tabel S7, see Fig. 6), suggesting that the effect the difference in drift rate of on response bias to /ti/ depends on the level of Chinese language proficiency. The simple slope analyses indicated that the difference of drift rate was positively related to the difference of response proportion among those who reported a high level of CLP (*B* = 0.239, *t* = 8.045, *p* < .001), not among those who reported a beginner-medium level of CLP (*B* = .033, *t* = −.459, *p* = .652). Individuals with high CLP exhibited a response bias pattern similar to that of native Chinese speakers, where syllable classification is influenced by speech comprehension. Specifically, the greater the difference in drift rates due to speech comprehension, the greater the difference in response bias on 50% morphing syllable. However, this is not observed in those with low CLP.

**Fig. 6.**
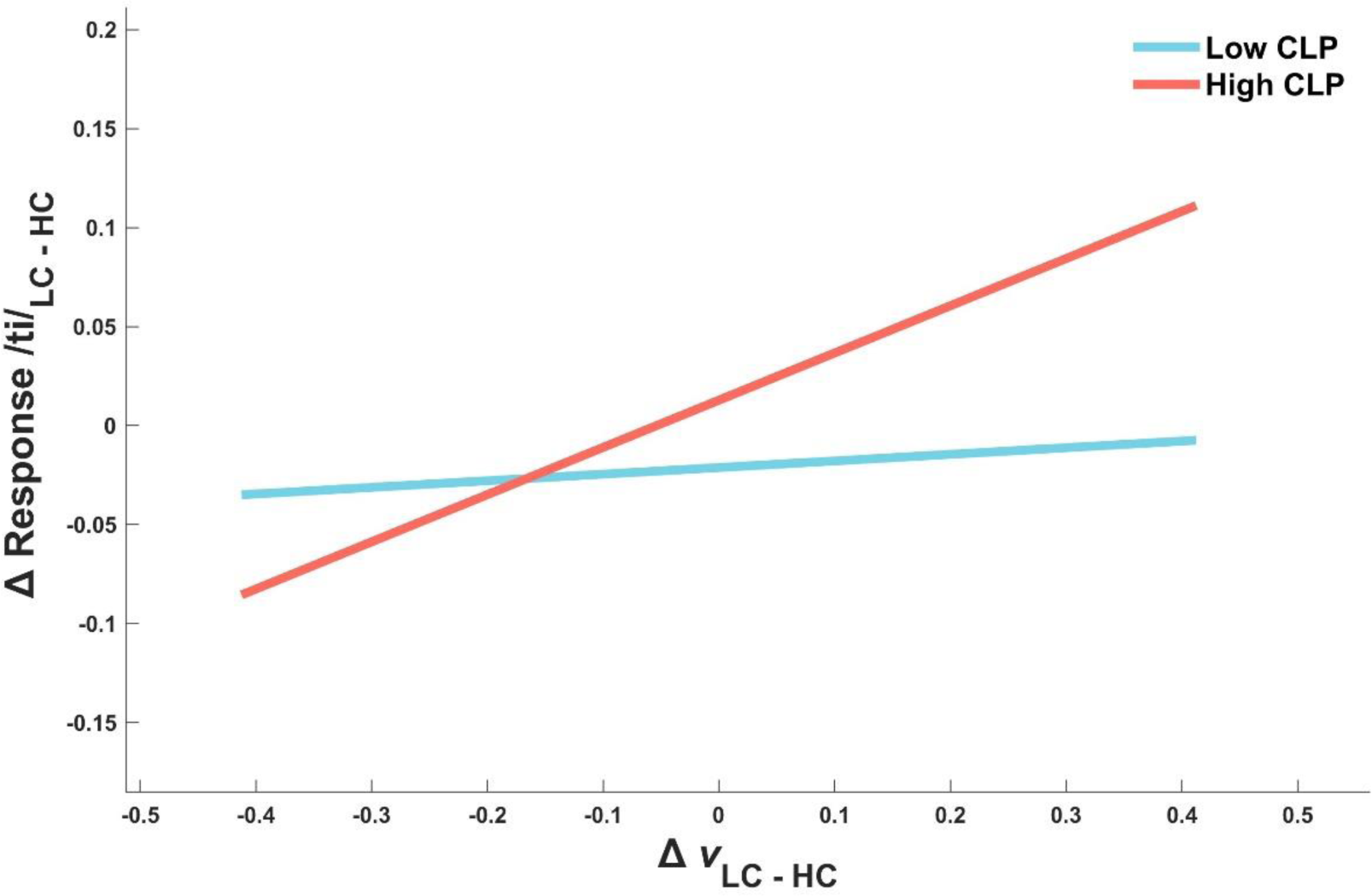
The simple slope analysis in 50% morphing level. *Note.* The difference of drift rate and the difference of responses proportion (/ti/) at different levels of CLP.

Surprisingly, in 25% and 75% morphing level, the CLP did not moderate the effect of the difference of drift rate on the response bias (25%: *B* = .008, *t* = .248, *p* = .807, *R^2^* change = .002, in Tabel S8; 75%: *B* = −.052, *t* = −1.282, *p* = .216, *R^2^* change = .038, in Tabel S9). We posit that in situations where the differences in subjective speech rate are less pronounced, listeners rely more on the distinctive features of the syllables for classification. The morphing syllables at the 25% and 75% levels provide more distinct features that aid in syllable classification, thus resulting in the absence of a moderation effect.

## Discussion

Time is relative. Our perception of time is a rather fragile mental construct, with changes in cognitive state or sensory environment making it appear to speed up or slow down. In Study 1, direct evidence showed that the acceleration of subjective speech rate on the low level of speech comprehension, relative to high level. Furthermore, we found in Study 2 that random sequences of syllables with low comprehensibility biased participants towards perceiving the consonant in a morphing syllable as /t/ (long voice onset time, VOT). Analysis using the drift diffusion model revealed this syllable classification bias occurred after syllable onset and could not be explained by the type of speech presented beforehand. Finally, in Study 3 among non-native Chinese speakers, we discovered that listener language proficiency modulated the impact of speech comprehension on subjective speech rate. Our research provides a novel perspective on studying time perception in complex stimuli by manipulating speech comprehension.

Our perceptual experiences are dynamic in nature. Not only do we experience time itself, but our experiences of time can speed up or slow down (Cohen, 2011; Kent & Wittmann, 2021). A key question is why merely changing speech comprehension leads listeners to perceive completely opposite speech rates. One potential explanation relates to event segmentation - how our brain parses continuous sensory inputs into discrete perceptual events to facilitate comprehension and memory (Bangert et al., 2020; Yousif & Scholl, 2019). When speech comprehension is reduced, it is difficult for listeners to segment speech streams into high-level semantic fragments. Instead, the original syllabic fragments are preserved (Blanco-Elorrieta et al., 2020; Ding et al., 2016). Listeners cannot segment or group low comprehensibility of speech into chunks of phrases or sentences, and more discrete syllables (or words in Chinese) are isolated as individual chunk, and less information is obtained from speech as temporal details are lost (Kurby & Zacks, 2008). This increases the overlapping boundaries between perceptual chunks. Distinguishing events within these overlapping boundaries is challenging, compressing the processed information, and leads to a shorter subjective duration (Bangert et al., 2020; Yousif & Scholl, 2019). Conversely, high comprehensibility of speech can be segmented into fewer and larger chunks, reducing the size of the overlapping area, and freeing up processing resources for temporal information. Therefore, it exhibits a longer perceived duration and shows that the speech rate is perceived as slower.

The findings from Study 3 provide further support for this event segmentation interpretation. We observed a moderating effect of language proficiency such that the drift rate could predict the bias of syllable classification in individuals with high Chinese proficiency, but not in those with low proficiency. Through language learning, non-native speakers can gradually grasp the boundaries between segmented words or sentences in that language, aiding the extraction of accurate semantic information (Chan et al., 2022; Yang, 2021). In contrast, individuals with low proficiency cannot effectively utilize grammatical rules, resulting in both speech types being perceived as discrete syllable segments without differences in subjective speech rate.

The perception of speech rate can occur at different timescales - from milliseconds at the word level, or seconds to minutes for whole speech passages. Time perception mechanisms differ between these scales. Faster, “automatic” processes underlie millisecond-range perception, while slower cognitive processes involving attention and memory dominate the supra-second range (Eagleman & Pariyadath, 2009; Grondin, 2010; Matthews & Meck, 2016). Our study does not determine whether the bias in speech rate perception stems from automatic or cognitive mechanisms. However, neurophysiological evidence allows some inferences. Previous research found greater prefrontal cortex activation during highly comprehensible speech processing (Lee et al., 2016), which is similar to activation seen in supra-second timing tasks (Basso et al., 2003; Lewis & Miall, 2003). In addition, higher trial-to-trial beta power indexes longer reproduced supra-second intervals in the supra-second interval timing task (Kononowicz & Rijn, 2015). Beta power can also reflect the comprehensibility of the speech being processed (Drijvers et al., 2018; Pefkou et al., 2017). Therefore, we hypothesize that the effect of speech comprehension on altering speech rate perception in our study may belong to supra-second cognitive time perception. But this interpretation requires further verification.

By the DDM, we did not find any differences in starting point between speech comprehension conditions. It ruled out the possibility that participants classified morphing syllables based on the preceding speech type. Conversely, when the upper boundary corresponded to syllable /ti/, the result showed that low comprehension produced a larger drift rate. Participants represent the preference for long VOT evidence accumulation (Ratcliff & McKoon, 2008). Furthermore, the drift criterion results demonstrated that speech comprehension altered the baseline of evidence collection during syllable processing (Osth et al., 2017). These findings indicate that the subjective speech rate bias induced by speech comprehension can produce the same rate normalization effect as objective speech rate differences (Bosker & Reinisch, 2017; Maslowski et al., 2019).

In summary, our findings suggest that changes in subjective speech rate may be driven by the speech comprehension. Unlike traditional time interval judgments, rate perception does not rely on accessing an internal representation of event onset and comparing it to the sensory information available at event offset to estimate how long the event lasted. It provides a window for probing time perception of complex stimuli. An outstanding question is how the precise scale of subjective speech rate interacts with language proficiency - this relationship could potentially serve as a novel indicator for second language assessment. More broadly, while humans are notoriously fickle at perceiving duration, studying time perception within speech processing sheds new light on the role of continuous perceptual encoding in shaping our experience of time. By bridging speech and time perception research, our work represents the first demonstration of how complex auditory inputs dynamically alters perceived rate.

## Supporting information

Sumpplementary

## Open Practices Statement

Three studies reported in this article were not preregistered before data collection. All data have been made publicly accessible at OSF and can be accessed at https://osf.io/bm6nx/. Analyzed codes are available upon request from the corresponding author. Our research was approved by the Committee for Protecting Human and Animal Subjects in School of Psychological and Cognitive Sciences at Peking University and the Institutional Review Board at Beijing Language and Culture University.

## Reference

Ahissar, E., Nagarajan, S., Ahissar, M., Protopapas, A., Mahncke, H., & Merzenich, M. M. (2001). Speech comprehension is correlated with temporal response patterns recorded from auditory cortex. Proceedings of the National Academy of Sciences, 98(23), 13367–13372. 10.1073/pnas.201400998

Baese-Berk, M. M., Heffner, C. C., Dilley, L. C., Pitt, M. A., Morrill, T. H., & McAuley, J. D. (2014). Long- Term Temporal Tracking of Speech Rate Affects Spoken-Word Recognition. Psychological Science, 25(8), 1546–1553. 10.1177/0956797614533705

Bangert, A. S., Kurby, C. A., Hughes, A. S., & Carrasco, O. (2020). Crossing event boundaries changes prospective perceptions of temporal length and proximity. *Attention, Perception*, & Psychophysics, 82(3), 1459–1472. 10.3758/s13414-019-01829-x

Basso, G., Nichelli, P., Wharton, C. M., Peterson, M., & Grafman, J. (2003). Distributed neural systems for temporal production: A functional MRI study. Brain Research Bulletin, 59(5), 405–411. 10.1016/S0361-9230(02)00941-3

Blanco-Elorrieta, E., Ding, N., Pylkkänen, L., & Poeppel, D. (2020). Understanding Requires Tracking: Noise and Knowledge Interact in Bilingual Comprehension. Journal of Cognitive Neuroscience, 32(10), 1975–1983. 10.1162/jocn_a_01610

Block, R. A., Hancock, P. A., & Zakay, D. (2010). How cognitive load affects duration judgments: A meta-analytic review. Acta Psychologica, 134(3), 330–343. 10.1016/j.actpsy.2010.03.006

Boersma, P. (2001). Praat, a system for doing phonetics by computer. Glot International, 5, 341–345.

Bosker, H. R., & Reinisch, E. (2017). Foreign Languages Sound Fast: Evidence from Implicit Rate Normalization. Frontiers in Psychology, 8, 1063. 10.3389/fpsyg.2017.01063

Chan, J., Woore, R., Molway, L., & Mutton, T. (2022). Learning and teaching Chinese as a foreign language: A scoping review. Review of Education, 10(3), e3370. 10.1002/rev3.3370

Cohen, M. X. (2011). It’s about Time. Frontiers in Human Neuroscience, 5. 10.3389/fnhum.2011.00002

Cui, X., Tian, Y., Zhang, L., Chen, Y., Bai, Y., Li, D., Liu, J., Gable, P., & Yin, H. (2023). The role of valence, arousal, stimulus type, and temporal paradigm in the effect of emotion on time perception: A meta-analysis. Psychonomic Bulletin & Review, 30(1), 1–21. 10.3758/s13423-022-02148-3

Ding, N., Melloni, L., Zhang, H., Tian, X., & Poeppel, D. (2016). Cortical tracking of hierarchical linguistic structures in connected speech. Nature Neuroscience, 19(1), 158–164. 10.1038/nn.4186

Drijvers, L., Özyürek, A., & Jensen, O. (2018). Hearing and seeing meaning in noise: Alpha, beta, and gamma oscillations predict gestural enhancement of degraded speech comprehension. Human Brain Mapping, 39(5), 2075–2087. 10.1002/hbm.23987

Eagleman, D. M. (2008). Human time perception and its illusions. Current Opinion in Neurobiology, 18(2), 131–136. 10.1016/j.conb.2008.06.002

Eagleman, D. M., & Pariyadath, V. (2009). Is subjective duration a signature of coding efficiency? Philosophical Transactions of the Royal Society B: Biological Sciences, 364(1525), 1841–1851. 10.1098/rstb.2009.0026

Einstein, A. (1920). Relativity: The Special and General Theory. Henry Holt and Company.

Eveline, V. (1982). Subjective Estimation of Speech Rate. Phonetica, 39(2–3), 136–149. 10.1159/000261656

Fernandes, A. C., & Garcia-Marques, T. (2020). A meta-analytical review of the familiarity temporal effect: Testing assumptions of the attentional and the fluency-attributional accounts. Psychological Bulletin, 146(3), 187–217. 10.1037/bul0000222

Grondin, S. (2010). Timing and time perception: A review of recent behavioral and neuroscience findings and theoretical directions. *Attention, Perception*, & Psychophysics, 72(3), 561–582. 10.3758/APP.72.3.561

Harvey, B. M., Dumoulin, S. O., Fracasso, A., & Paul, J. M. (2020). A Network of Topographic Maps in Human Association Cortex Hierarchically Transforms Visual Timing-Selective Responses. Current Biology, 30(8), 1424–1434.e6. 10.1016/j.cub.2020.01.090

Hayes, A. (2013). Introduction to mediation, moderation, and conditional process analysis: A regression-based approach. The Guilford Press.

Howard, M. W. (2018). Memory as Perception of the Past: Compressed Time in Mind and Brain. Trends in Cognitive Sciences, 22(2), 124–136. 10.1016/j.tics.2017.11.004

Jacoby, L. L., & Dallas, M. (1981). On the Relationship Between Autobiographical Memory and Perceptual Learning. Journal of Experimental Psychology. General, 110(3), 306–340. 10.1037/0096-3445.110.3.306

JASP Team (2023). JASP (Version 0. 17.3) [Computer software].

Kent, L., & Wittmann, M. (2021). Time consciousness: The missing link in theories of consciousness. Neuroscience of Consciousness, 2021(2), niab011. 10.1093/nc/niab011

Kloosterman, N. A., De Gee, J. W., Werkle-Bergner, M., Lindenberger, U., Garrett, D. D., & Fahrenfort, J. J. (2019). Humans strategically shift decision bias by flexibly adjusting sensory evidence accumulation. ELife, 8, e37321. 10.7554/eLife.37321

Kononowicz, T. W., & Rijn, H. van. (2015). Single trial beta oscillations index time estimation. Neuropsychologia, 75, 381–389. 10.1016/j.neuropsychologia.2015.06.014

Kurby, C. A., & Zacks, J. M. (2008). Segmentation in the perception and memory of events. Trends in Cognitive Sciences, 12(2), 72–79. 10.1016/j.tics.2007.11.004

Lee, Y.-S., Min, N. E., Wingfield, A., Grossman, M., & Peelle, J. E. (2016). Acoustic richness modulates the neural networks supporting intelligible speech processing. Hearing Research, 333, 108–117. 10.1016/j.heares.2015.12.008

Lewis, P. A., & Miall, R. C. (2003). Distinct systems for automatic and cognitively controlled time measurement: Evidence from neuroimaging. Current Opinion in Neurobiology, 13(2), 250–255. 10.1016/S0959-4388(03)00036-9

Maslowski, M., Meyer, A. S., & Bosker, H. R. (2019). Listeners normalize speech for contextual speech rate even without an explicit recognition task. The Journal of the Acoustical Society of America, 146(1), 179–188. 10.1121/1.5116004

Matthews, W. J., & Meck, W. H. (2016). Temporal cognition: Connecting subjective time to perception, attention, and memory. Psychological Bulletin, 142(8), 865–907. 10.1037/bul0000045

McGettigan, C., Faulkner, A., Altarelli, I., Obleser, J., Baverstock, H., & Scott, S. K. (2012). Speech comprehension aided by multiple modalities: Behavioural and neural interactions. Neuropsychologia, 50(5), 762–776. 10.1016/j.neuropsychologia.2012.01.010

Moore, B. C. J. (2019). The roles of temporal envelope and fine structure information in auditory perception. Acoustical Science and Technology, 40(2), 61–83. 10.1250/ast.40.61

Naghibi, N., Jahangiri, N., Khosrowabadi, R., Eickhoff, C. R., Eickhoff, S. B., Coull, J. T., & Tahmasian, M. (2023). Embodying Time in the Brain: A Multi-Dimensional Neuroimaging Meta-Analysis of 95 Duration Processing Studies. Neuropsychology Review. 10.1007/s11065-023-09588-1

Osth, A. F., Dennis, S., & Heathcote, A. (2017). Likelihood ratio sequential sampling models of recognition memory. Cognitive Psychology, 92, 101–126. 10.1016/j.cogpsych.2016.11.007

Pefkou, M., Arnal, L. H., Fontolan, L., & Giraud, A.-L. (2017). θ-Band and β-Band Neural Activity Reflects Independent Syllable Tracking and Comprehension of Time-Compressed Speech. The Journal of Neuroscience, 37(33), 7930–7938. 10.1523/JNEUROSCI.2882-16.2017

Pfitzinger, H. R., & Tamashima, M. (2006). Comparing Perceptual Local Speech Rate of German and Japanese Speech. 105–108. http://www.isca-speech.org/archive.

Plug, L., & Smith, R. (2021). The role of segment rate in speech tempo perception by English listeners. Journal of Phonetics, 86, 101040. 10.1016/j.wocn.2021.101040

Price, C. (2003). An Overview of Speech Comprehension and Production. In Human Brain Function (Second Edition) (pp. 517–532). Academic Press.

Ratcliff, R., & McKoon, G. (2008). The Diffusion Decision Model: Theory and Data for Two-Choice Decision Tasks. Neural Computation, 20(4), 873–922. 10.1162/neco.2008.12-06-420

Ratcliff, R., & Rouder, J. N. (1998). Modeling Response Times for Two-Choice Decisions. Psychological Science, 9(5), 347–356. 10.1111/1467-9280.00067

Ratcliff, R., Sederberg, P. B., Smith, T. A., & Childers, R. (2016). A single trial analysis of EEG in recognition memory: Tracking the neural correlates of memory strength. Neuropsychologia, 93, 128–141. 10.1016/j.neuropsychologia.2016.09.026

Swaminathan, J., Mason, C. R., Streeter, T. M., Best, V., Roverud, E., & Kidd, G. (2016). Role of Binaural Temporal Fine Structure and Envelope Cues in Cocktail-Party Listening. Journal of Neuroscience, 36(31), 8250–8257. 10.1523/JNEUROSCI.4421-15.2016

Wade, T., & Holt, L. L. (2005). Perceptual effects of preceding nonspeech rate on temporal properties of speech categories. Perception & Psychophysics, 67(6), 939–950. 10.3758/BF03193621

Wang, J., Chen, J., Yang, X., Liu, L., Wu, C., Lu, L., Li, L., & Wu, Y. (2021). Common Brain Substrates Underlying Auditory Speech Priming and Perceived Spatial Separation. Frontiers in Neuroscience, 15, 664985. 10.3389/fnins.2021.664985

Wänke, M., & Hansen, J. (2015). Relative Processing Fluency. Current Directions in Psychological Science, 24(3), 195–199. 10.1177/0963721414561766

Wiecki, T. V., Sofer, I., & Frank, M. J. (2013). HDDM: Hierarchical Bayesian estimation of the Drift-Diffusion Model in Python. Frontiers in Neuroinformatics, 7. 10.3389/fninf.2013.00014

Winn, M. B. (2020). Manipulation of voice onset time in speech stimuli: A tutorial and flexible Praat script. The Journal of the Acoustical Society of America, 147(2), 852–866. 10.1121/10.0000692

Yang, S. (2021). Investigating word segmentation of chinese second language learners. Reading and Writing, 34(5), 1273–1293. 10.1007/s11145-020-10113-6

Yousif, S. R., & Scholl, B. J. (2019). The one-is-more illusion: Sets of discrete objects appear less extended than equivalent continuous entities in both space and time. Cognition, 185, 121–130. 10.1016/j.cognition.2018.10.002

