## Supplementary material for "The More Unclear the Hearing, the Faster the Feeling: Speech Comprehension Changes Subjective Speech Rate": Sumpplementary

**Study 1: Listeners judge the speech rate in noise-vocoder speech.**

**Method**

***Clear Speech****.* Speaker was asked to keep neutral emotions and read naturally and at a constant rate. Noise reduction was carried out on the recording to eliminate potential ambient noise as much as possible. In addition, irrelevant acoustic fragments such as exhalation at the beginning of each sentence were manually retrieved and deleted, and 25 milliseconds at the onset and offset of each sentence were smoothed through the cosine window. Spectral analysis was performed for the time domain envelope of all the recordings. The speech envelopes showed a strong oscillatory component at 3.7 Hz (see Fig. S1). The rate corresponds to the syllabic presentation rate of the stimuli. For visualization, the speech envelope power spectra have been normalized by dividing the power spectra by their maximum power value.

**Fig. S1.**

*Speech envelope power spectra (average across all sentences).*

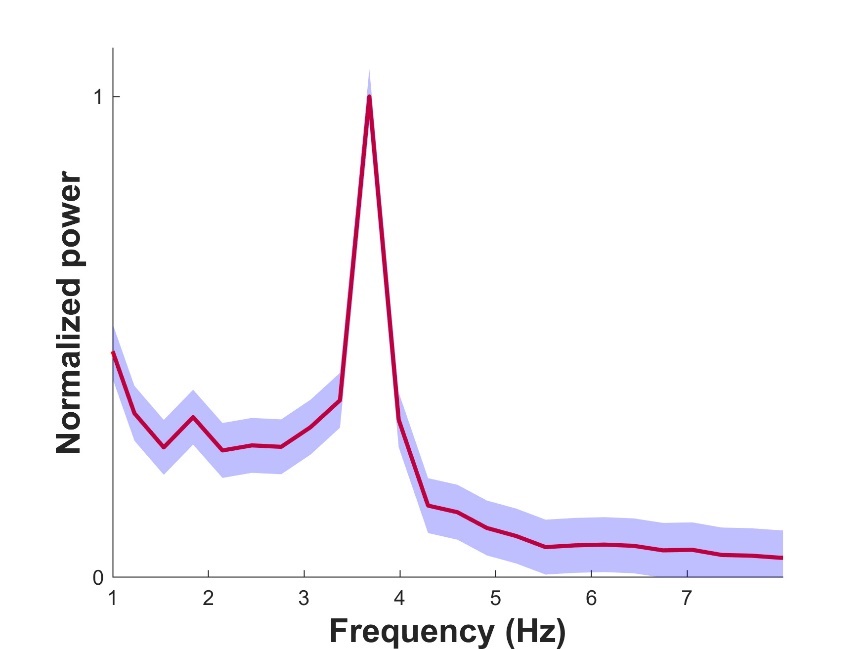

*Note.* The blue areas represent standard errors.

We recruited 50 Native Chinese speakers to rating all speech. The speech rate is scored from 1 to 5, with 1 being very slow and 5 being very fast. Similarly, the understanding score also ranges from 1 to 5, with 1 being difficult to understand and 5 being easy to understand. All ratings data were analyzed by computing the interclass correlation coefficient (ICC2k; two-way random model, absolute agreement) alongside their 95% confidence intervals (CIs) as a measure of interrater agreement using the *psych* package in R (version 4.3.1). The ICC ranges between 0 (no agreement) and 1 (perfect agreement). In these scores, high agreement indicated that listeners agreed with each other on which speech similar on the speech rate and understanding. Conversely, low agreement was seen as evidence that likely no consistency was formed and that listeners provided noisy/random ratings. In the Speech rate score, the ICC2k was .808, and the mean range of all scores was from 3.14 to 4.56 (see Fig. S2 left). In the understanding score, the ICC2k was .858, and the mean range of all scores was from 2.8 to 3.8 (see Fig. S2 right). Taken together, the substantial agreement across listeners shows that for speech rate and understanding, there is no obvious difference among all the speech.

**Fig. S2.**

*The result of interclass correlation coefficient in speech rate and understanding rating.*

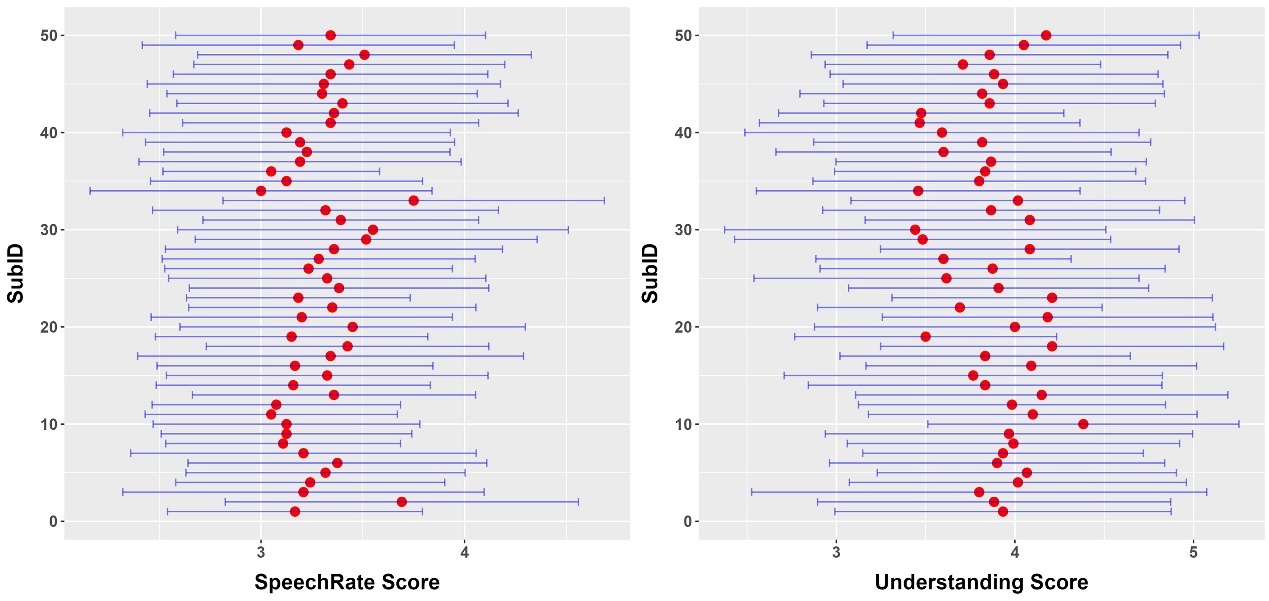

*Note.* The left graph shows the score of speech rate of all evaluators, and the right graph shows the score of understanding. The red dot represents the average score of all sentences, and the blue error bar is the standard deviation.

***Noise-vocoder speech*.** We first filtered the natural speech and a speech-spectrum noise (same power spectrum as the natural speech) into different bands. According to behavioral evidence (Newman & Chatterjee, 2013), noise-vocoded speech with 2 band is unintelligible. In order to find the understanding threshold of close to 50% for each participant, we created a total of 2-8 bands of all speech. The frequency range is from 80 to 8820Hz and is spaced in equal steps along the cochlear frequency map according to the number of frequency bands (Greenwood, 1990). Secondly, we extracted the envelope and the temporal fine structure (TFS) from the filtered signals of both the natural speech and a speech-spectrum noise. Thirdly, the envelope of each filter output from the natural speech was then multiplied by the TFS of the corresponding filter output from the speech-spectrum noise (Smith et al., 2002), resulting in natural speech-noise chimeras. Finally, we summed these chimeras (with energy normalized) to produce a noise vocoded speech (NV).

**Results**

***The accuracy of speech comprehension test****.* Figure S3 presents the accuracy of verbal repeating of speech with HC and LC condition. Through paired sample t-test, it was found that the keyword accuracy of speech with high speech comprehension is higher than that of speech with low speech comprehension (*t* = 7.791, *df* = 30, *p* < .001, Cohen’s *d* = 1.399; HC: *M* = .806, *SE* = .016; LC: *M* = .562, *SE* = .031). This indicates that the speech familiarity task indeed improves the comprehensibility of noise-vocoder speech.

**Fig. S3.**

*The accuracy of speech comprehension test between high speech comprehension and low speech comprehension condition.*

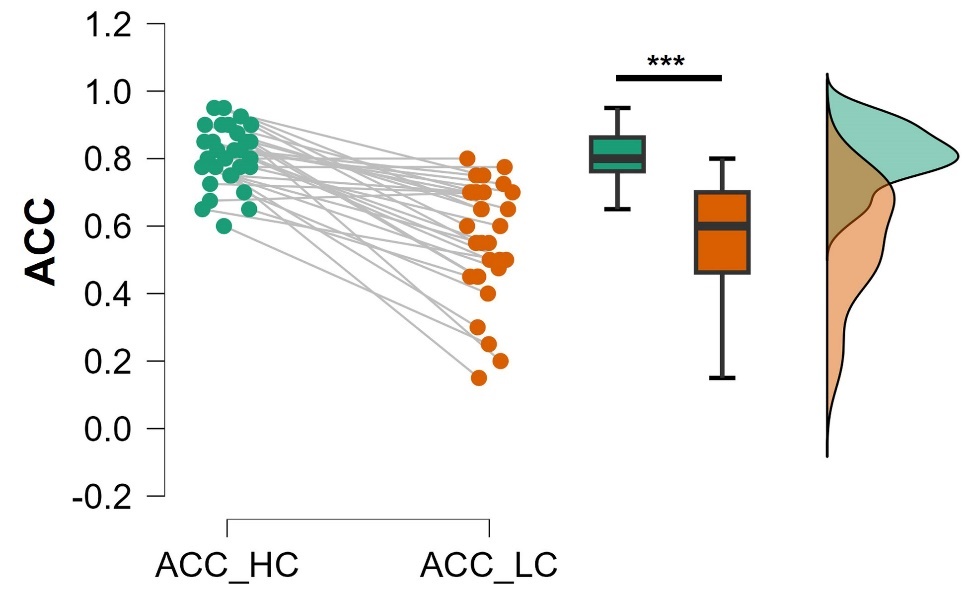

*Note.* Solid dots represent data for each subject. ***: *p* < .001.

***The PSE of high and low comprehension condition.*** Figure S4 presents the PSE values under the HC and LC conditions respectively. The results of the single-sample t-test show that the PSE of both high comprehensibility (*M* = .978, *SE* = .004; *t* = -4.990, *df* = 30, *p* < .001, Cohen’s *d* = -.896) and low comprehensibility (*M* = .988, *SE* = .005; *t* = -2.487, *df* = 30, *p* = .019, Cohen’s *d* = -.447) are less than the standard speech rate (i.e., PSE = 1).

**Fig. S4.**

*The PSE between high speech comprehension and low speech comprehension condition.*

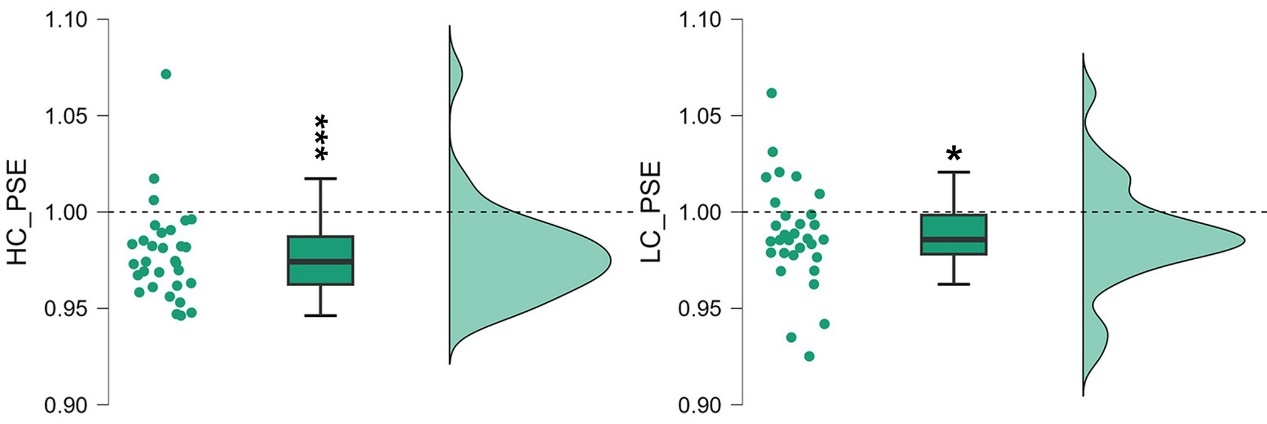

*Note.* Solid dots represent data for each subject. ***: *p* < .001; *: *p* < .05

**Study 2: Subjective speech rate was assessed by implicit speech rate task.**

**Method**

***Stimuli.*** All syllables of simple sentences and random syllable sequences were synthesized independently using the speech synthesizer (https://www.readspeaker.com/, the male speaker, Liang). The durations of synthesized syllable were 83-334 ms (mean duration 208 ms). Then all syllables were adjusted to 250 ms by truncation or padding silence at the end with a 25-ms cosine-squared falling ramp. No acoustic gaps were inserted between syllables. Thus, each speech, which of the speech rate was fixed at 4 Hz, was an isochronous sequence of syllables. Three speeches of the same type of sentence will be randomly selected for each trial, which was 3 seconds.

The morphing syllables is /di/-/ti/ continuum. We progressively deleted the onset of a syllable (/ti/) with a voiced stop sound and replaced it with a roughly equivalent amount of the onset from its voiceless onset counterpart. In order to exclude the influence of fundamental frequency (F0) cues on syllable classification, F0 was controlled to be equalized across all members of morphing syllables. All operations are carried out through the open-source software Praat (Boersma & Weenink, 2023). These stimuli were normed for the root-mean-square intensity to broadly equalize the perceived loudness. As in study 1, all stimuli were always presented binaurally via headphone.

**Results**

***DDM general parameters.***

**Table S1.**

*Summary statistics for each group level parameter’s posterior in target model*

|  | mean | std | 2.5q | 25q | 50q | 75q | 97.5q | mc err |
| --- | --- | --- | --- | --- | --- | --- | --- | --- |
| a | 1.521 | 0.048 | 1.424 | 1.490 | 1.521 | 1.551 | 1.617 | 0.001 |
| a_std | 0.243 | 0.038 | 0.180 | 0.216 | 0.239 | 0.265 | 0.328 | 0.001 |
| v(HC.25%) | -1.020 | 0.169 | -1.350 | -1.132 | -1.021 | -0.908 | -0.684 | 0.003 |
| v(HC.50%) | 0.368 | 0.171 | 0.044 | 0.251 | 0.366 | 0.483 | 0.709 | 0.003 |
| v(HC.75%) | 1.806 | 0.171 | 1.476 | 1.690 | 1.806 | 1.919 | 2.144 | 0.004 |
| v(LC.25%) | -0.591 | 0.167 | -0.922 | -0.700 | -0.594 | -0.475 | -0.265 | 0.003 |
| v(LC.50%) | 0.574 | 0.168 | 0.245 | 0.460 | 0.573 | 0.688 | 0.895 | 0.003 |
| v(LC.75%) | 1.918 | 0.171 | 1.581 | 1.803 | 1.918 | 2.037 | 2.247 | 0.003 |
| v_std | 0.832 | 0.054 | 0.733 | 0.794 | 0.830 | 0.868 | 0.945 | 0.001 |
| t | 0.207 | 0.080 | 0.029 | 0.156 | 0.221 | 0.267 | 0.330 | 0.002 |
| t_std | 0.234 | 0.059 | 0.147 | 0.190 | 0.224 | 0.270 | 0.373 | 0.002 |
| z(HC) | 0.507 | 0.009 | 0.489 | 0.501 | 0.507 | 0.513 | 0.525 | 0.000 |
| z(LC) | 0.510 | 0.009 | 0.491 | 0.504 | 0.510 | 0.516 | 0.529 | 0.000 |
| z_std | 0.126 | 0.030 | 0.068 | 0.105 | 0.125 | 0.146 | 0.188 | 0.002 |

***DDM posterior predictive check.***

**Table S2.**

*Summary statistics for each of the simulated data sets from the posterior and the observed data in target model.*

| State | M_obs | M_stim | std | SEM | MSE | credible | quantile | mahalanobis |
| --- | --- | --- | --- | --- | --- | --- | --- | --- |
| P_resp | .619 | .619 | .319 | .000 | .102 | TRUE | 43.301 | .000 |
| M_ub | .723 | .773 | .278 | .003 | .080 | TRUE | 46.452 | 0.181 |
| sd_ub | .421 | 0.344 | 0.194 | 0.006 | 0.043 | TRUE | 72.072 | 0.398 |
| 10q_ub | .283 | 0.457 | 0.196 | 0.030 | 0.069 | TRUE | 29.425 | 0.890 |
| 30q_ub | .497 | 0.556 | 0.217 | 0.004 | 0.050 | TRUE | 39.410 | 0.274 |
| 50q_ub | .640 | 0.676 | 0.253 | 0.001 | 0.065 | TRUE | 45.388 | 0.143 |
| 70q_ub | .815 | 0.851 | 0.319 | 0.001 | 0.103 | TRUE | 50.035 | 0.112 |
| 90q_ub | 1.254 | 1.195 | 0.482 | 0.003 | 0.236 | TRUE | 61.144 | 0.122 |
| M_lb | -.794 | -0.797 | 0.289 | 0.000 | 0.084 | TRUE | 46.851 | 0.011 |
| sd_lb | .430 | 0.329 | 0.220 | 0.010 | 0.059 | TRUE | 72.815 | 0.460 |
| 10q_lb | .363 | 0.496 | 0.212 | 0.018 | 0.063 | TRUE | 31.191 | 0.630 |
| 30q_lb | .571 | 0.592 | 0.228 | 0.000 | 0.053 | TRUE | 44.519 | 0.094 |
| 50q_lb | 0.700 | 0.708 | 0.264 | 0.000 | 0.070 | TRUE | 50.910 | 0.030 |
| 70q_lb | 0.869 | 0.875 | 0.334 | 0.000 | 0.111 | TRUE | 54.080 | 0.018 |
| 90q_lb | 1.318 | 1.190 | 0.513 | 0.016 | 0.279 | TRUE | 65.630 | 0.248 |

**Fig. S5**.

*Posterior predictive check by group in target model*

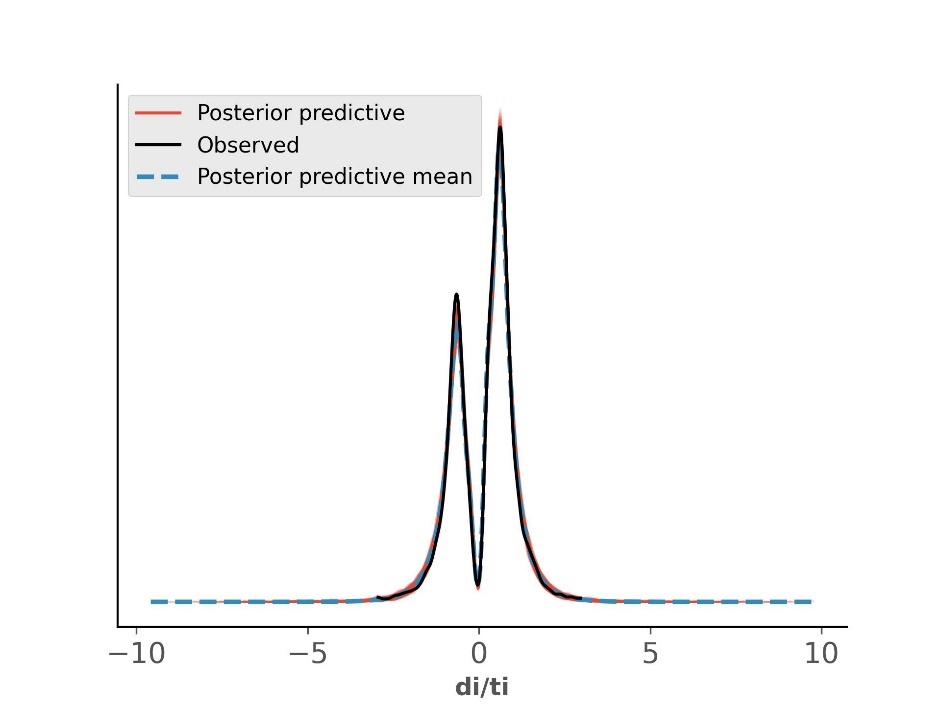

*The drift criterion model.* Individual’s response bias may be caused by perceptual processing, characterized by drift rate (Ratcliff & McKoon, 2008; Ratcliff & Rouder, 1998). In the analysis of drift rate, we have found that the drift rate of participants in syllable classification is not the same under different speech comprehension conditions. This bias can be conceptualized as a drift criterion (dc), which is the amount added/subtracted on an unbiased drift rate (Ratcliff & McKoon, 2008). Specifically, the drift criterion is equivalent to adding a constant unrelated to evidence in the drift, so a non-zero drift criterion leads to a linear increase in decision variable bias over time (see Fig. S6). It should be noted that to estimate dc, we must set the correct answer for each syllable classification, so we exclude the 50% level of morphing syllable level. For the 25% level, we stipulate that choosing /di/ is the correct answer. For the 75%, we stipulate that choosing /ti/ is the correct answer. The null model is set to all parameters being fixed values between conditions, and the target model is set to dc changing with speech comprehension conditions.

**Fig. S6.**

*The drift criterion diagram.*

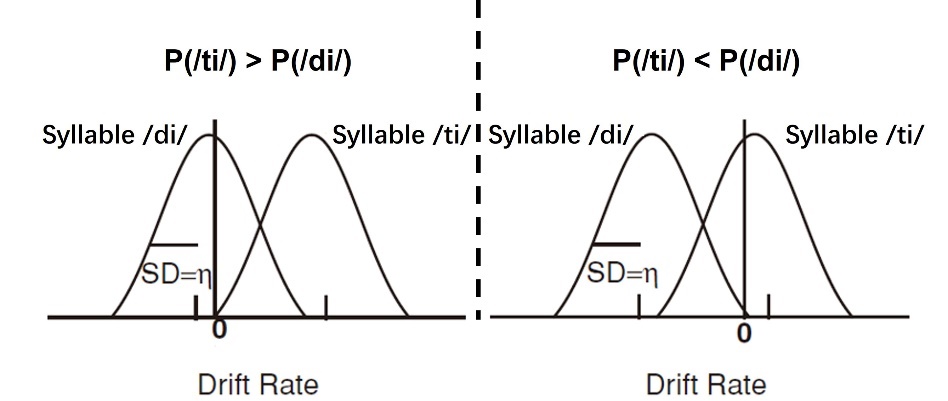

*Note*. drift criterion (the zero point) varying with probability. When the probability of response /ti/ is higher, the zero point tend to the left side of the evidence distribution. This is similar to the criterion in signal detection theory that only a little evidence can be detected or accumulated.

The null model was set so that all parameters are fixed across conditions. The dc parameters of the target model are freely estimated with the conditions of speech comprehension, and the other parameters are fixed between the conditions. We also used MCMC sampling to estimate the posterior distribution of each model and ran 4 chains with 5000 samples, with the first 500 samples discarded as burn-in. Convergence was assessed by visually inspection (the trace and autocorrelation) and the Gelman-Rubin R̂ statistic. The range of R̂ statistic values across all group parameter in each model was 1 to 1.01, suggesting good convergence. Goodness of fit was visually inspected with a posterior predictive check (see Fig. S7).

**Fig. S7**.

*Posterior predictive check by group in target model (dc).*

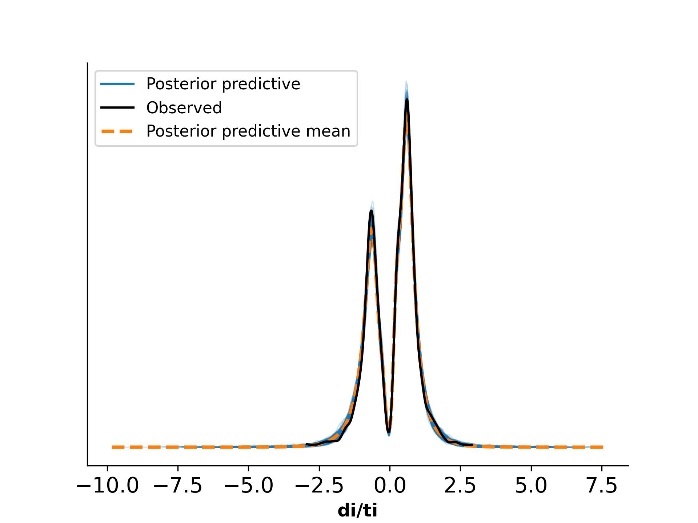

The results of the model comparison show that the DIC of the target model is lower (delta DIC relative to null model: = -36.407), indicating that the change in dc can explain more behavioral data. Figure S8 presents the posterior probability distribution of dc under different conditions, and it can be found that dc is larger under the LC condition (LC>HC: posterior *p* = .968). This verifies the conclusion that the bias of syllable classification is due to the preference for evidence selection.

**Fig. S8.**

*Posterior probability distribution of parameter dc.*

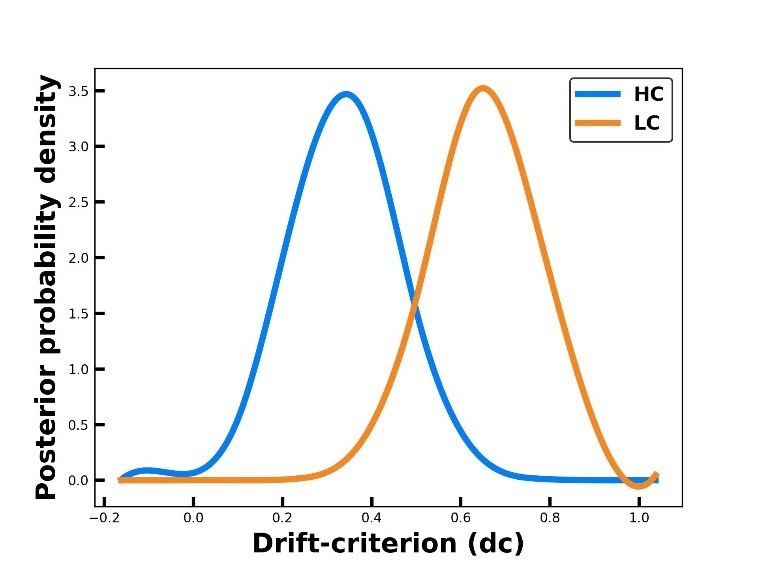

**Table S3**

*Summary statistics for each group level parameter’s posterior in target model(dc).*

|  | mean | std | 2.5q | 25q | 50q | 75q | 97.5q | mc err |
| --- | --- | --- | --- | --- | --- | --- | --- | --- |
| a | 1.525 | 0.051 | 1.428 | 1.491 | 1.525 | 1.557 | 1.628 | 0.001 |
| a_std | 0.242 | 0.039 | 0.176 | 0.214 | 0.237 | 0.265 | 0.330 | 0.001 |
| v | 1.328 | 0.157 | 1.017 | 1.226 | 1.326 | 1.432 | 1.641 | 0.002 |
| v_std | 0.794 | 0.125 | 0.595 | 0.706 | 0.777 | 0.867 | 1.068 | 0.002 |
| t | 0.307 | 0.042 | 0.235 | 0.278 | 0.303 | 0.331 | 0.400 | 0.001 |
| t_std | 0.216 | 0.044 | 0.150 | 0.185 | 0.209 | 0.241 | 0.319 | 0.001 |
| z | 0.517 | 0.007 | 0.502 | 0.513 | 0.517 | 0.522 | 0.532 | 0.000 |
| z_std | 0.086 | 0.030 | 0.018 | 0.068 | 0.088 | 0.107 | 0.142 | 0.002 |
| dc(HC) | 0.349 | 0.110 | 0.127 | 0.277 | 0.349 | 0.421 | 0.566 | 0.002 |
| dc(LC) | 0.631 | 0.109 | 0.415 | 0.560 | 0.631 | 0.703 | 0.847 | 0.002 |
| dc_std | 0.524 | 0.063 | 0.413 | 0.480 | 0.520 | 0.562 | 0.660 | 0.001 |

**Table S4.**

*Summary statistics for each of the simulated data sets from the posterior and the observed data in target model (dc).*

| State | M_obs | M_stim | std | SEM | MSE | credible | quantile | mahalanobis |
| --- | --- | --- | --- | --- | --- | --- | --- | --- |
| P_resp | 0.603 | 0.595 | 0.364 | 0.000 | 0.133 | TRUE | 47.220 | 0.022 |
| M_ub | 0.688 | 0.729 | 0.268 | 0.002 | 0.073 | TRUE | 47.838 | 0.153 |
| sd_ub | 0.399 | 0.303 | 0.186 | 0.009 | 0.044 | TRUE | 76.025 | 0.513 |
| 10q_ub | 0.266 | 0.451 | 0.199 | 0.034 | 0.074 | TRUE | 27.735 | 0.927 |
| 30q_ub | 0.479 | 0.539 | 0.215 | 0.004 | 0.050 | TRUE | 39.186 | 0.277 |
| 50q_ub | 0.620 | 0.644 | 0.247 | 0.001 | 0.062 | TRUE | 46.804 | 0.099 |
| 70q_ub | 0.774 | 0.798 | 0.306 | 0.001 | 0.094 | TRUE | 52.759 | 0.079 |
| 90q_ub | 1.164 | 1.098 | 0.453 | 0.004 | 0.210 | TRUE | 63.664 | 0.147 |
| M_lb | -0.762 | -0.770 | 0.287 | 0.000 | 0.083 | TRUE | 47.334 | 0.030 |
| sd_lb | 0.411 | 0.295 | 0.215 | 0.013 | 0.059 | TRUE | 74.566 | 0.539 |
| 10q_lb | 0.352 | 0.500 | 0.225 | 0.022 | 0.073 | TRUE | 30.573 | 0.659 |
| 30q_lb | 0.554 | 0.588 | 0.236 | 0.001 | 0.057 | TRUE | 43.084 | 0.145 |
| 50q_lb | 0.679 | 0.692 | 0.266 | 0.000 | 0.071 | TRUE | 50.348 | 0.049 |
| 70q_lb | 0.828 | 0.842 | 0.328 | 0.000 | 0.108 | TRUE | 53.179 | 0.042 |
| 90q_lb | 1.253 | 1.124 | 0.498 | 0.016 | 0.265 | TRUE | 66.444 | 0.258 |

**Study 3: Language proficiency moderates the effect of speech comprehension on implicit speech rate.**

**Results**

***DDM general parameters****.*

**Table S5**.

*Summary statistics for each group level parameter’s posterior in target model (non-native Chinses speakers’ group).*

|  | mean | std | 2.5q | 25q | 50q | 75q | 97.5q | mc err |
| --- | --- | --- | --- | --- | --- | --- | --- | --- |
| a | 1.478 | 0.041 | 1.395 | 1.451 | 1.477 | 1.504 | 1.559 | 0.001 |
| a_std | 0.200 | 0.034 | 0.146 | 0.175 | 0.196 | 0.221 | 0.278 | 0.001 |
| v(HC.25%) | -1.133 | 0.157 | -1.441 | -1.239 | -1.133 | -1.031 | -0.824 | 0.003 |
| v(HC.50%) | -0.221 | 0.154 | -0.521 | -0.325 | -0.221 | -0.116 | 0.073 | 0.002 |
| v(HC.75%) | 0.848 | 0.157 | 0.536 | 0.744 | 0.849 | 0.957 | 1.157 | 0.003 |
| v(LC.25%) | -1.154 | 0.157 | -1.465 | -1.261 | -1.151 | -1.045 | -0.850 | 0.003 |
| v(LC.50%) | -0.204 | 0.152 | -0.505 | -0.308 | -0.203 | -0.099 | 0.093 | 0.003 |
| v(LC.75%) | 0.844 | 0.160 | 0.529 | 0.735 | 0.844 | 0.949 | 1.158 | 0.003 |
| v_std | 0.724 | 0.051 | 0.629 | 0.689 | 0.722 | 0.757 | 0.829 | 0.001 |
| t | 0.518 | 0.013 | 0.494 | 0.510 | 0.518 | 0.526 | 0.543 | 0.000 |
| t_std | 0.061 | 0.010 | 0.045 | 0.054 | 0.060 | 0.067 | 0.084 | 0.000 |
| z(HC) | 0.526 | 0.012 | 0.501 | 0.518 | 0.526 | 0.534 | 0.550 | 0.000 |
| z(LC) | 0.526 | 0.013 | 0.502 | 0.518 | 0.526 | 0.535 | 0.551 | 0.000 |
| z_std | 0.196 | 0.032 | 0.139 | 0.173 | 0.194 | 0.216 | 0.264 | 0.001 |

***DDM posterior predictive check****.*

**Table S6.**

*Summary statistics for each of the simulated data sets from the posterior and the observed data in target model (non-native Chinses speakers’ group).*

| State | M_obs | M_stim | std | SEM | MSE | credible | quantile | mahalanobis |
| --- | --- | --- | --- | --- | --- | --- | --- | --- |
| P_resp | 0.476 | 0.465 | 0.297 | 0.000 | 0.088 | TRUE | 54.824 | 0.037 |
| M_ub | 0.946 | 0.996 | 0.212 | 0.002 | 0.047 | TRUE | 44.918 | 0.234 |
| sd_ub | 0.359 | 0.341 | 0.180 | 0.000 | 0.033 | TRUE | 59.488 | 0.102 |
| 10q_ub | 0.612 | 0.687 | 0.135 | 0.006 | 0.024 | TRUE | 25.734 | 0.560 |
| 30q_ub | 0.734 | 0.781 | 0.156 | 0.002 | 0.027 | TRUE | 42.758 | 0.304 |
| 50q_ub | 0.853 | 0.898 | 0.193 | 0.002 | 0.039 | TRUE | 45.958 | 0.234 |
| 70q_ub | 1.009 | 1.071 | 0.256 | 0.004 | 0.069 | TRUE | 45.292 | 0.241 |
| 90q_ub | 1.410 | 1.407 | 0.409 | 0.000 | 0.168 | TRUE | 55.524 | 0.007 |
| M_lb | -1.001 | -1.033 | 0.204 | 0.001 | 0.043 | TRUE | 53.658 | 0.158 |
| sd_lb | 0.350 | 0.354 | 0.174 | 0.000 | 0.030 | TRUE | 53.458 | 0.024 |
| 10q_lb | 0.649 | 0.706 | 0.117 | 0.003 | 0.017 | TRUE | 29.131 | 0.489 |
| 30q_lb | 0.786 | 0.810 | 0.142 | 0.001 | 0.021 | TRUE | 48.321 | 0.171 |
| 50q_lb | 0.917 | 0.935 | 0.182 | 0.000 | 0.033 | TRUE | 50.983 | 0.100 |
| 70q_lb | 1.099 | 1.115 | 0.248 | 0.000 | 0.062 | TRUE | 51.753 | 0.065 |
| 90q_lb | 1.466 | 1.466 | 0.403 | 0.000 | 0.162 | TRUE | 54.328 | 0.001 |

**Figure S9**.

*Posterior predictive check by group in target model (non-native Chinses speakers’ group).*

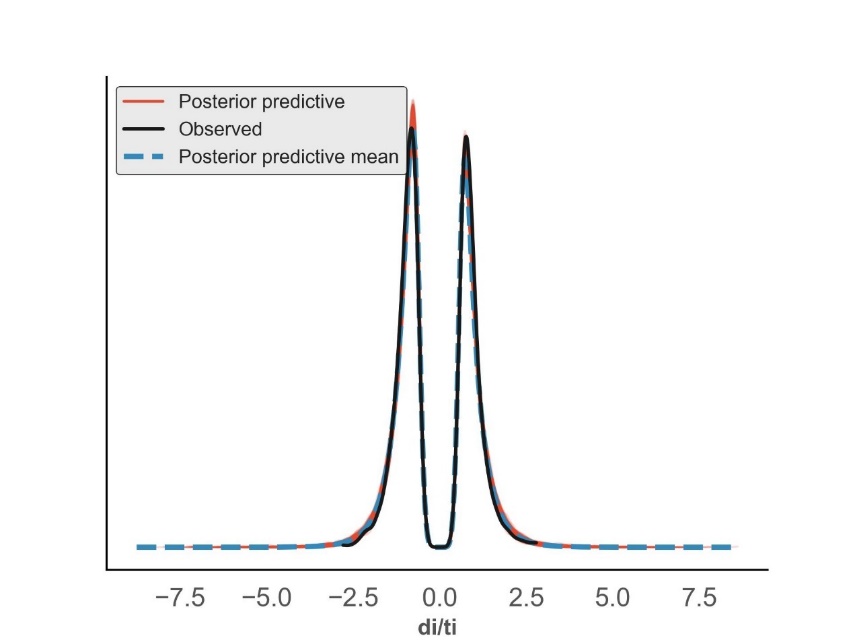

***The moderating effect models.*** The moderation analysis examined the potential conditional effects of Chinese language proficiency on the direct process between the drift rate and the bias classification. Findings in the 50% level morphing syllable is shown in Table S7.

**Table S7.**

*Ordinary least squares regression results for the moderated model in 50% morphing syllable condition.*

| **Variables** | *B* | *SE* | *t* | *p* | *LLCI* | *ULCI* |
| --- | --- | --- | --- | --- | --- | --- |
| Outcomes: △response proportion for /ti/ | | | | | | |
| Covariates | | | | | | |
| Age | -.0014 | .0032 | -.4269 | .6743 | -.0082 | .0054 |
| Gender | -.0336 | .0207 | -1.6191 | .1219 | -.0769 | .0098 |
| Predictors | | | | | | |
| △*v* | .1359 | .0383 | 3.5526 | .0021** | .0558 | .2160 |
| CLP | .0170 | .0097 | 1.7609 | .0943 | -.0032 | .0372 |
| △*v* * CLP | .1026 | .0401 | 2.5621 | .0191* | .0188 | .1865 |

*Note*. Bootstrap sample size = 5,000. LLCI = lower limit confidence interval 95 %. ULCI = upper limit confidence interval 95 %. *v* = the drift rate. CLP = Chinese language proficiency. ⁎⁎*p* < .01, ⁎*p* < .05.

Findings in the 25% level morphing syllable is shown in Table S8.

**Table S8**.

*Ordinary least squares regression results for the moderated model in 25% morphing syllable condition.*

| **Variables** | *B* | *SE* | *t* | *p* | *LLCI* | *ULCI* |
| --- | --- | --- | --- | --- | --- | --- |
| Outcomes: △response proportion for /ti/ | | | | | | |
| Covariates | | | | | | |
| Age | -.0020 | .0036 | -.5554 | .5851 | -.0094 | .0055 |
| Gender | -.0177 | .0241 | -.7322 | .4730 | -.0682 | .0328 |
| Predictors | | | | | | |
| △*v* | .0726 | .0320 | 2.2679 | .0352* | .0056 | .1396 |
| CLP | .0112 | .0107 | 1.0505 | .3064 | -.0112 | .0336 |
| △*v* * CLP | .0078 | .0314 | 0.2478 | .8069 | -.0579 | .0734 |

*Note*. Bootstrap sample size = 5,000. LLCI = lower limit confidence interval 95 %. ULCI = upper limit confidence interval 95 %. *v* = the drift rate. CLP = Chinese language proficiency. ⁎*p* < .05.

Findings in the 75% level morphing syllable is shown in Table S9.

**Table S9.**

*Ordinary least squares regression results for the moderated model in 75% morphing syllable condition.*

| **Variables** | *B* | *SE* | *t* | *p* | *LLCI* | *ULCI* |
| --- | --- | --- | --- | --- | --- | --- |
| Outcomes: △response proportion for /ti/ | | | | | | |
| Covariates | | | | | | |
| Age | .0027 | .0040 | .6682 | .5120 | -.0057 | .0111 |
| Gender | -.0388 | .0275 | -1.4116 | .1742 | -.0965 | .0188 |
| Predictors | | | | | | |
| △*v* | .1624 | .0393 | 4.1360 | .0006*** | .0802 | .2447 |
| CLP | .0039 | .0125 | 0.3095 | .7603 | -.0222 | .0299 |
| △*v* * CLP | -.0507 | .0396 | -1.2807 | .2157 | -.1337 | .0322 |

*Note*. Bootstrap sample size = 5,000. LLCI = lower limit confidence interval 95 %. ULCI = upper limit confidence interval 95 %. *v* = the drift rate. CLP = Chinese language proficiency. *** *p* < .001.

**Reference**

Boersma, Paul & Weenink, David (2023). Praat: doing phonetics by computer [Computer program]. Version 6.3.13, retrieved 31 July 2023 from <http://www.praat.org/>

Greenwood, D. D. (1990). A cochlear frequency-position function for several species—29 years later. *The Journal of the Acoustical Society of America*, *87*(6), 2592–2605. <https://doi.org/10.1121/1.399052>

Newman, R., & Chatterjee, M. (2013). Toddlers’ recognition of noise-vocoded speech. *The Journal of the Acoustical Society of America*, *133*(1), 483–494. <https://doi.org/10.1121/1.4770241>

Ratcliff, R., & McKoon, G. (2008). The Diffusion Decision Model: Theory and Data for Two-Choice Decision Tasks. *Neural Computation*, *20*(4), 873–922. https://doi.org/10.1162/neco.2008.12-06-420

Ratcliff, R., & Rouder, J. N. (1998). Modeling Response Times for Two-Choice Decisions. *Psychological Science*, *9*(5), 347–356. https://doi.org/10.1111/1467-9280.00067

Smith, Z. M., Delgutte, B., & Oxenham, A. J. (2002). Chimaeric sounds reveal dichotomies in auditory perception. *Nature*, *416*(6876), 87–90. https://doi.org/10.1038/416087a
